# Accurate Predictions of Liquid-Liquid Phase Separating Proteins at Single Amino Acid Resolution

**DOI:** 10.1101/2024.07.19.602785

**Authors:** Michele Monti, Jonathan Fiorentino, Dimitrios Miltiadis-Vrachnos, Giorgio Bini, Tiziana Cotrufo, Natalia Sanchez de Groot, Alexandros Armaos, Gian Gaetano Tartaglia

## Abstract

Liquid-liquid phase separation (LLPS) is a molecular mechanism that leads to the formation of membraneless organelles inside the cell. Despite recent advances in the experimental probing and computational prediction of proteins involved in this process, the identification of the protein regions driving LLPS and the prediction of the effect of mutations on LLPS are lagging behind.

Here, we introduce catGRANULE 2.0 ROBOT (R - Ribonucleoprotein, O - Organization, in B - Biocondensates, O - Organelle, T - Types), an advanced algorithm for predicting protein LLPS at single amino acid resolution. Integrating physico-chemical properties of the proteins and structural features derived from AlphaFold models, catGRANULE 2.0 ROBOT significantly surpasses traditional sequence-based and state-of-the-art structure-based methods in performance, achieving an Area Under the Receiver Operating Characteristic Curve (AUROC) of 0.76 or higher. We present a comprehensive evaluation of the algorithm across multiple organisms and cellular components, demonstrating its effectiveness in predicting LLPS propensities at the single amino acid level and the impacts of mutations on LLPS. Our results are robustly supported by experimental validations, including immunofluorescence microscopy images from the Human Protein Atlas.

catGRANULE 2.0 ROBOT’s potential in protein design and mutation control can improve our understanding of proteins’ propensity to form subcellular compartments and help develop strategies to influence biological processes through LLPS. catGRANULE 2.0 ROBOT is freely available at https://tools.tartaglialab. com/catgranule2.

## 1 Introduction

Liquid-liquid phase separation (LLPS) is a molecular phenomenon that brings molecules together to form membraneless condensates inside the cell [6, 36, 41, 42, 55]. Recent studies evidenced the key role of LLPS in human health and disease, especially in protein condensation and neurodegenerative disorders [37]. RNA is known to play a central role in LLPS; indeed, proteins undergo phase separation only in specific transcriptomic conditions [21, 79]. Moreover, condensates formed through phase separation can catalyze crucial biochemical processes by concentrating and compartmentalizing specific proteins at precise subcellular locations [5].

Contrary to the process of liquid to solid phase transition (LSPT), in which proteins go toward an irreversible aggregation state [54, 80], LLPS is a reversible process [50]. The reversibility of LLPS has a dual function: while the increase in protein concentration enhances the enzymatic activity [20, 44, 79], RNA accumulation in organelles such as P-bodies can inhibit protein translation [34].

Despite recent advances in the experimental probing and characterization of LLPS [3] and the construction of large-scale databases of LLPS-prone proteins [39, 43, 53, 65, 68, 75], a wide coverage of the proteome of different species in light of their LLPS properties is still lacking. For this reason, several computational methods have been developed to predict the propensity of a protein to undergo LLPS [7, 23, 30, 47, 59]. However, most of these methods lack the ability to predict the LLPS propensity at the amino acid level and none of them has been extensively tested for the prediction of the effect of single and multiple mutations of the protein sequence on their capability to undergo LLPS. One of the first LLPS predictors, catGRANULE 1.0, computes the protein propensity for granule formation based on structural disorder and nucleic acid-binding propensities [7]. Following this, the MaGS method was developed using a variety of features including protein abundance, intrinsic disorder percentage, phosphorylation site annotations, PScore, Camsol score, RNA interaction, and the composition of leucine and glycine [30, 31]. A more recent method is PICNIC, which uses both sequence-based and structure-based features derived from AlphaFold2 models, focusing on sequence complexity, disorder scores, and amino acid co-occurrences [23]. An extended version, PICNICGO, adds Gene Ontology terms to provide deeper insights into functions like RNA-binding [23]. Lastly, PSPHunter broadens the feature set further by implementing word2vec for sequence analysis, alongside Position-Specific Scoring Matrix (PSSM) and Hidden Markov Model (HMM) to capture evolutionary and structural insights, encompassing a wide array of functional traits like protein modifications and network properties [59].

In this work, we introduce an advanced predictor of LLPS proteins called catGRAN-ULE 2.0 ROBOT (R - Ribonucleoprotein, O - Organization, in B - Biocondensates, O - Organelle, T - Types). This new version significantly enhances the capabilities of its predecessor, catGRANULE 1.0, and it is based on a curated database of phase-separating proteins and their mutants. catGRANULE 2.0 ROBOT integrates a comprehensive set of features that include structural and sequence-based data derived from AlphaFold2 models, specifically targeting properties relevant to phase separation. This version has undergone testing against a wide array of mutations, which were meticulously compiled through an exhaustive literature search. Unlike PSPHunter, catGRANULE 2.0 ROBOT opts out of using encoders of protein sequence features, choosing instead to merge predictions based on physico-chemical properties derived from sequence and structural data. This approach is designed to enhance the interpretability of the model’s predictions. With the continuous expansion in the number of known phase-separating proteins, catGRANULE 2.0 ROBOT takes an approach to balance between interpretability and performance, aiming to provide a robust tool to study protein phase separation.

The extensive training dataset used in catGRANULE 2.0 ROBOT comprises human proteins documented to undergo LLPS, sourced from various databases and resources [31, 39, 43, 53, 65, 68, 75, 76]. It also includes a selection of negative proteins—those highly unlikely to undergo phase separation, specifically excluding known interactors of LLPS proteins [45]. In this manuscript, we first report the characterization of proteins belonging to the training dataset compared to the rest of the human proteome [40, 51]. Following this, we describe each protein in the dataset using a list of 128 features that considers both sequence-based physico-chemical properties of the protein and structural properties, which we extract based on the AlphaFold Structure Database [27, 64]. After we encode sequence- and structure-based features into a vector, we train multiple binary classifiers and we select the best performing model, which we call ROBOT, to define a LLPS propensity score for a protein. We tested catGRANULE 2.0 ROBOT on proteins belonging to different condensates in human and in different organisms and we show that it outperforms previous methods, both based on sequence features [7, 30, 31, 59] but also on structural features [23]. Moreover, we provide an orthogonal validation of our predictions using thousands of antibody-based immunofluorescence (IF) confocal microscopy images obtained from the Human Protein Atlas [63].

catGRANULE 2.0 ROBOT designs profiles of LLPS propensity along protein sequences and accurately identifies regions experimentally confirmed as LLPS drivers. Furthermore, catGRANULE 2.0 ROBOT can assess the impact of single and multiple amino acid mutations on LLPS propensity, determining whether mutations will increase or decrease it. To this end, we use mutations identified through a comprehensive literature review, including a deep mutational scanning of TDP-43 [8].

Finally, we developed a user-friendly web server (https://tools.tartaglialab.com/ catgranule2) to make our algorithm easily usable by the scientific community. In conclusion, we introduced a method that outperforms previous algorithms in predicting the likelihood of proteins undergoing LLPS, computes a profile at the amino acid level, accurately predicts the effect of mutations, and can be used to design proteins and mutations with adjustable LLPS properties.

## 2 Results

### 2.1 Construction and biological characterization of the training dataset

With the aim of building a robust machine learning method to predict the LLPS propensity of proteins at the amino acid level, we defined training and test datasets with the following workflow. We first collected human proteins known to be involved in LLPS from several publicly available databases [31, 43, 53, 65, 68, 75, 76], obtaining 5656 LLPS-prone proteins in total (Figure 1A-B; see Methods). We built the negative set by removing these proteins and their first interactors from the human proteome (see Methods) [45]. To prevent overfitting during the training phase, we utilized CD-HIT [17] to filter both positive and negative sets, ensuring sequence similarity was below 50%. Subsequently, we divided the data into training and test sets, as detailed in Supplementary Figure S1A-B (refer to Methods section for more information).

**Fig. 1.**
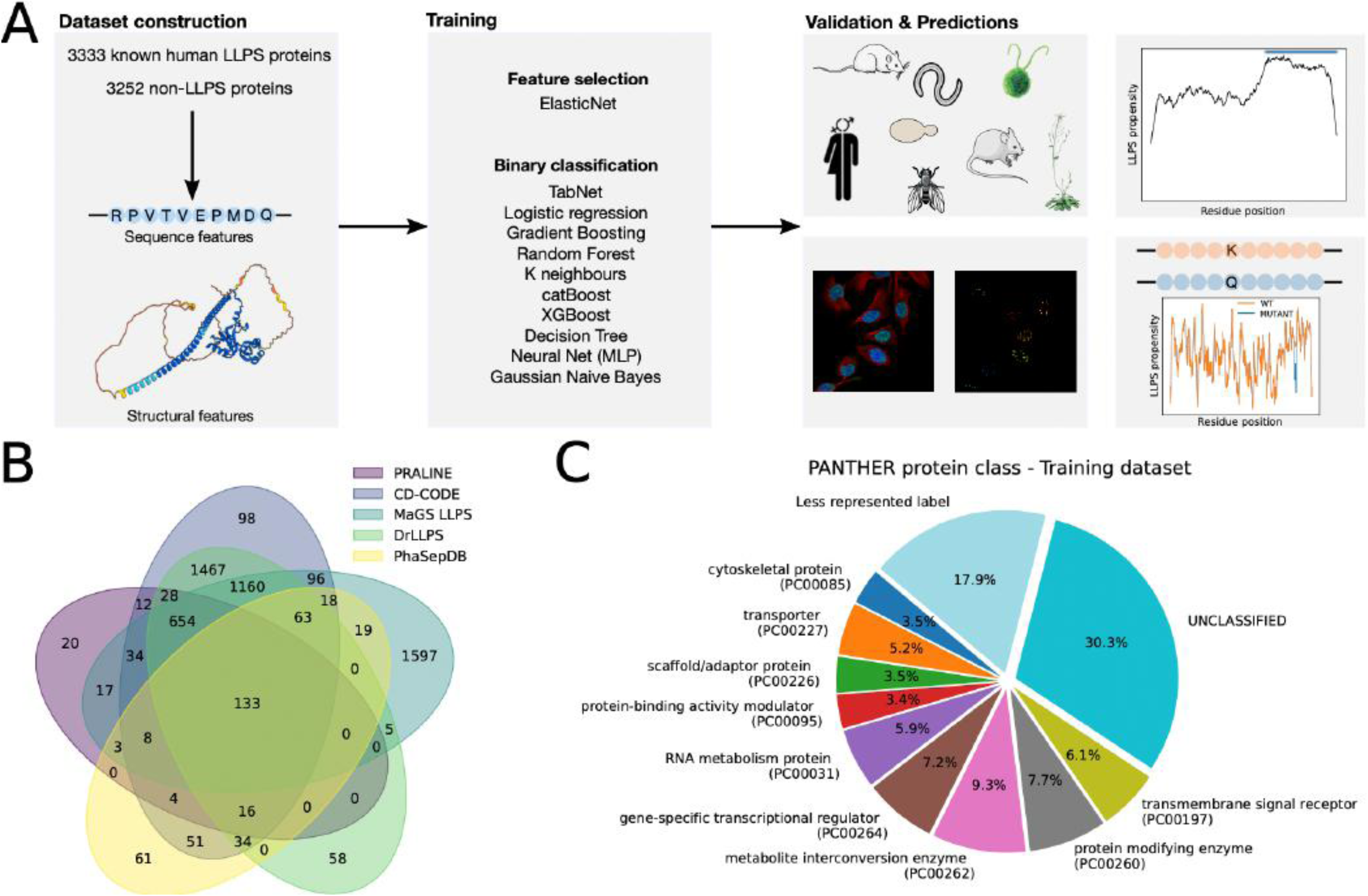
A. Schematics of the catGRANULE 2.0 ROBOT workflow. A training dataset is constructed consisting of 3333 known human LLPS proteins and 3252 non-LLPS proteins. The proteins are then encoded in a set of 128 features, including sequence-based physico-chemical and Alphafold2-derived structural features. Next, a subset of rel_2_ev_9_ant features is selected using ElasticNet and ten different classifiers are trained on the dataset; MLP is the selected classifier according to its superior performance on the test dataset. catGRANULE 2.0 ROBOT predictions are then validated on sets of known LLPS-prone proteins from different species [43] and on immunofluorescence microscopy images from the Human Protein Atlas. LLPS propensity profiles are predicted with a sliding window approach and validated on experimentally known LLPS driving regions of proteins belonging to different species, obtained from the PhaSepDB database [75]. Finally, catGRANULE 2.0 ROBOT predicts the effect of single and multiple amino acid mutations on LLPS propensity. B. Venn diagram showing the overlap of LLPS-prone proteins collected from different databases. C. Composition of the training dataset in terms of Panther protein class categories. Protein classes with less than 3 % have been aggregated in the ”Less represented label” category.

Comparing the distribution of the length and abundance of the LLPS proteins from the training set with the negative set (Supplementary Figure S1C-D), we observe that the differences are consistent with a comparison against the rest of the human proteome (Supplementary Figure S1E-F).

Next, we performed a Gene Ontology (GO) term enrichment analysis using Panther [40], which revealed that the LLPS proteins in our training set are significantly enriched in protein classes linked to RNA-related activities, translation, protein binding, and metabolic processes (Figure 1C and Supplementary Figure S2), compared to the rest of the proteome, while the proteins in the negative set are enriched for transporters and transmembrane receptors proteins (Supplementary Figure S2). These findings are consistent with the literature. RNA metabolism proteins such as TIA-1 and G3BP1 facilitate stress granule formation through LLPS, emphasizing the role of RNA-binding proteins in cellular stress responses [52]. Protein modifying enzymes influence LLPS via various post-translational modifications that alter protein interactions and stability [32]. In contrast, transporter and membrane proteins often form irreversible aggregates [61]. Additionally, defense and immunity proteins tend to undergo LSPT, leading to protein aggregates associated with diseases like ALS, indicating a distinct behavior in protein phase transitions [19]; see Methods and Supplementary Table S1. Although we found that proteins belonging to the positive and negative training sets are enriched in different biological features, we do not observe a strong separation of the two sets from a Principal Components Analysis (PCA) (Supplementary Figure S1G), motivating us to rely on non linear machine learning methods for the classification task.

We characterized each protein in the dataset with a set of 128 features, reported in Supplementary Table S2. We incorporated 80 physico-chemical features derived from protein sequence analysis and 2 phenomenological sequence patterns (see Methods) [7, 29], along with 28 structural features extracted using predicted protein structures from the AlphaFold Structure Database [27, 64]. This approach enabled us to identify features related to both the surface and the inner parts of the protein. Additionally, we incorporated 18 features based on the compositional similarity of protein sequence windows to experimentally determined RNA-binding patches, enhancing our ability to identify potential RNA-interacting regions in the protein sequences [13] (see Methods). In Supplementary Figure S3 we show a cluster map of the correlation matrix between the set of 128 features for the training dataset. We observe the presence of large clusters, especially for subsets of the physico-chemical features, as expected given the higher redundancy in their collection, compared to the structural features (see Methods).

### 2.2 catGRANULE 2.0 ROBOT accurately classifies LLPS prone proteins

After the construction of robust training and test datasets, we developed a machine learning pipeline to predict the LLPS propensity of a protein. We employed ElasticNet [81] to identify the most relevant features for LLPS classification, then we trained 10 different binary classifiers, performing a grid search over the hyper-parameters of each classifier and employing 5-fold Cross Validation during training to avoid overfitting (see Methods). Finally, we tested their performance on an independent test dataset using the Area Under the Receiver-Operating Characteristic Curve (AUROC) as scoring metric.

We found that the trained classifiers yield comparable AUROC scores and we selected the Multi-Layer Perceptron (MLP) as the optimal one, based on its superior performance on the test dataset (Supplementary Figure S4A). We compared the performance of our trained model on the test dataset with catGRANULE 1.0 [7], and the top performing state-of-the-art methods that are MaGS [30, 31], PICNIC, PICNIC-GO [23] and PSPHunter [59] (see Methods), observing that our model (catGRANULE 2.0 ROBOT) emerges as the best one (Figure 2A), although with an AUROC score slightly higher than the one of the other methods. Additionally, we found that catGRANULE 2.0 ROBOT outperforms the other predictors using other performance metrics, such as the accuracy, the F1-score, the Matthew’s correlation coefficient (MCC) and the recall, while MaGS, PSPHunter and PICNIC-GO perform better only for the precision (Supplementary Figure S4B). We provide the LLPS score computed using catGRAN-ULE 2.0 ROBOT for the whole human proteome in Supplementary Table S3. As an additional validation of the good performances of our model, we tested its capability to predict the LLPS propensity of proteins other than human, which was the only organism considered during the training (Figure 2B, Supplementary Figure S4C and Supplementary Table S4). In Figure 2B we show the fraction of correctly predicted proteins for several species, and we compare the result with PICNIC when available [23]. We observe that catGRANULE 2.0 ROBOT significantly outperforms PICNIC in most of the species considered and it performs comparably in the others.

**Fig. 2.**
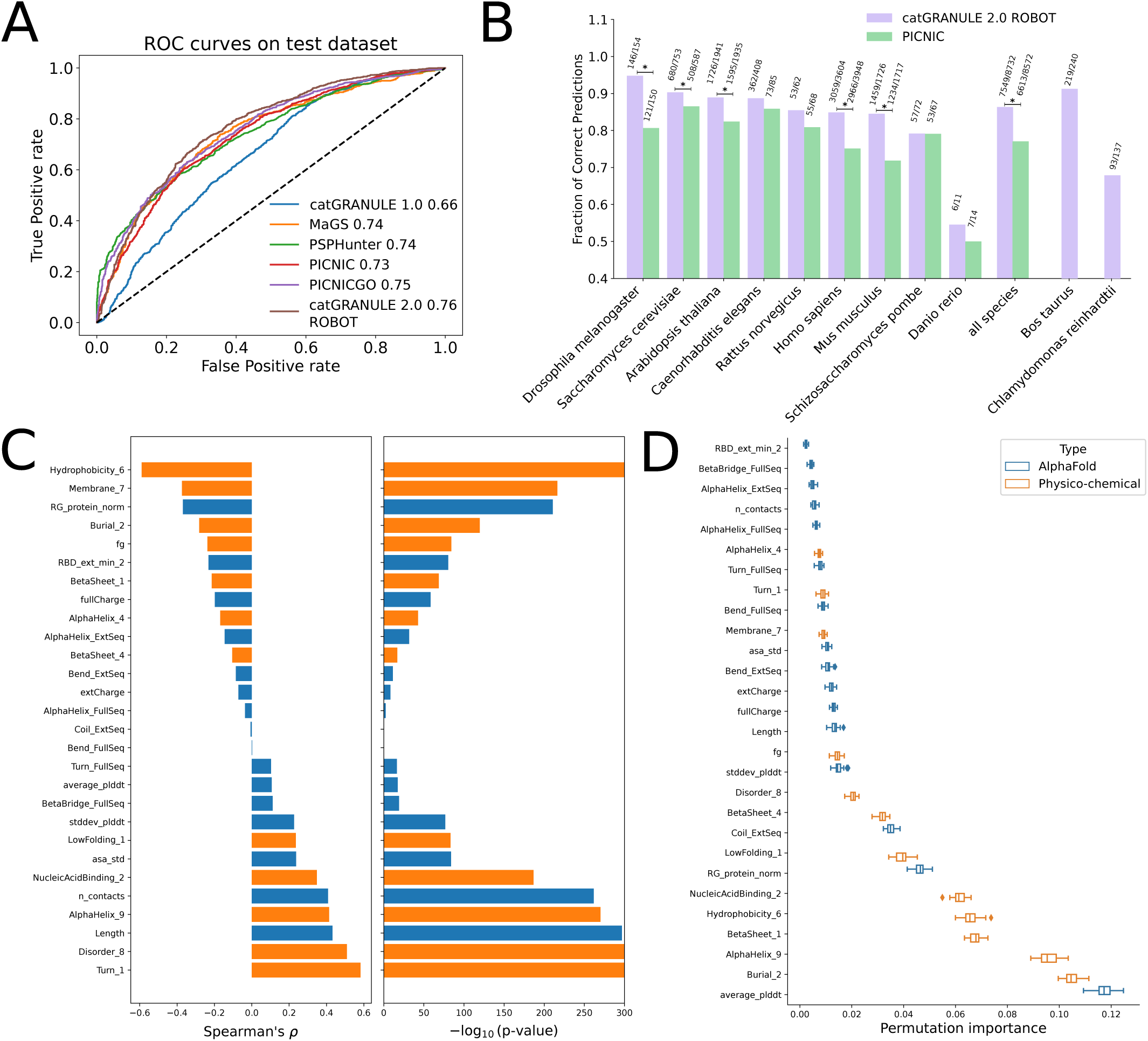
A. Receiver-Operating Characteristic (ROC) curves obtained from the test dataset, for catGRANULE 2.0 ROBOT and other LLPS prediction algorithms (see Methods for details). The Area under the ROC curve (AUROC) for each algorithm is indicated in the legend. B. Bar plot of the fraction of correctly predicted LLPS proteins for different species. The annotation of LLPS proteins was obtained from the DrLLPS database [43]. A star above a bar indicates a p-value smaller than 0.05 from a Fisher’s exact test between the fraction of correctly predicted LLPS proteins in catGRANULE 2.0 ROBOT and in PICNIC. C. Bar plot showing the Spearman’s correlation coefficient between the 28 features selected during the training step using ElasticNet and the predicted LLPS score, for the proteins belonging to the training dataset. The bar plot on the right shows the -log10(p-value) of the correlation coefficient. D. Box plot of the permutation importance computed on the training dataset for the 28 features selected during the training step using ElasticNet.

Using ElasticNet, we identified 28 features that are the most relevant to distinguish LLPS proteins from the negative set. Interestingly, while some of these features strongly separate the two classes individually, others do not show such strong differences between the classes, justifying our choice of a multivariate and non linear feature selection method (Supplementary Figure S5A and see Methods). This result was further corroborated by a comparison of the performances obtained with the 28 features selected by ElasticNet with those achieved by a linear model trained on all the 128 features, or with a MLP model trained adding features iteratively using a univariate feature selection method (Supplementary Figure S5B; see Methods).

Afterwards, to quantify the impact of the features to the discrimination of LLPS-prone proteins, we computed the Spearman’s correlation coefficient of each selected feature with the predicted LLPS score, using the proteins belonging to the training set (Figure 2C). We found, as expected and in agreement with the results from the catGRANULE 1.0 algorithm [7], that features associated to hydrophobicity (e.g., Hydrophobicity 6) are negatively associated with the LLPS propensity score [3], while those related to the nucleic acid binding propensity (e.g., NucleicAcidBinding 2) [79], to the disorder (e.g., Disorder 8) [26] or to the protein length [7] are positively associated. We found that the radius of gyration normalized by protein length (RG protein norm) exhibits a negative correlation with the LLPS score, while the standard deviation of the Accessible Surface Area (asa std) shows a positive correlation (Figure 2C). This suggests that more compact proteins are less prone to liquid-liquid phase separation (LLPS). Indeed, the power-law exponent *<* 1.0 for soluble species (monomers and oligomers, [60]) suggest that phase-separating proteins display consistent scaling behavior in terms of size and shape, a fundamental characteristic of protein structure. So, proteins with variable surface exposure, which facilitates interactions with other proteins and nucleic acids, are more likely to undergo LLPS. Coiled coils deviate from this typical behavior by exhibiting a linear scaling of the radius of gyration with the number of residues, further emphasizing the influence of specific protein structures on LLPS propensity. Then, we computed the permutation importance to quantify the contribution of each feature to the overall score (Figure 2D; see Methods). We found that the average pLDDT is the most relevant feature. This finding aligns with expectations, as pLDDT is a measure of protein disorder [71], which corroborates our analysis that protein disorder significantly contributes to phase separation [7].

We further characterized the significance and the hierarchy of the selected features by recursively eliminating the features selected by ElasticNet, then repeating the feature selection and training steps of our model (Supplementary Figure S5C; see Methods). We observe that the first large drop in performance happens after 6 iterations (Supplementary Figure S5C), when the size of the feature pool is approximately a quarter of the original one and the most informative feature families (e.g., Burial, BetaSheet, AlphaHelix, Geometry) are almost extinguished (Supplementary Figure S5D). Once the most important features are removed, the properties selected are of the same kind but are less effective as predictors. This consistency in the type of features chosen, even after primary ones are excluded, indicates that the method is stable with respect to the selection of properties, further validating our feature selection process. For instance, after the first iteration, the feature NucleicAcidBinding 2 is substituted by NucleicAcidBinding 7, while features related to the alpha helix (both sequence-based, AlphaHelix 4 and AlphaHelix 9, and structure-based, AlphaHelix FullSeq and AlphaHelix ExtSeq) are subsituted by AlphaHelix 1, AlphaHelix 6 and AlphaHelix 7 (Supplementary Table S2). These findings are significant, underscoring the importance of properties such as nucleic acid binding in LLPS. Specifically, these nucleic acid binding predictions are based on the electrostatic charge, which is known to facilitate RNA contact [21, 54] and prevent protein aggregation [11].

As a further validation of catGRANULE 2.0 ROBOT predictions, we used 10757 antibody-based images obtained by immunofluorescence (IF) confocal microscopy in human cell lines obtained from the Human Protein Atlas (https://www.proteinatlas.org/humanproteome/cell) [63]. After cell segmentation, we computed the coefficient of variation (CV) of the green fluorescence per cell and we considered the maximum of this quantity over the cells, for each protein. Next, we identified puncta of the green fluorescent protein and we computed the area, normalized by the average area of the nuclei per image, and the average number of puncta per protein (see Methods and Supplementary Table S5). We chose these quantities since we hypothesized that proteins undergoing LLPS would have more and larger droplets compared to other proteins, and a more compartmentalized expression. Indeed, ranking the proteins according to the LLPS propensity score predicted by catGRANULE 2.0 ROBOT and considering an increasing number of top and bottom proteins, we found that the AUROC scores start from values ranging between 0.6 and 0.7, depending on the quantity under study, and decrease toward 0.5, as shown in Figure 3A, demonstrating that proteins with higher predicted LLPS score show droplets with the hypothesized features in IF images.

**Fig. 3.**
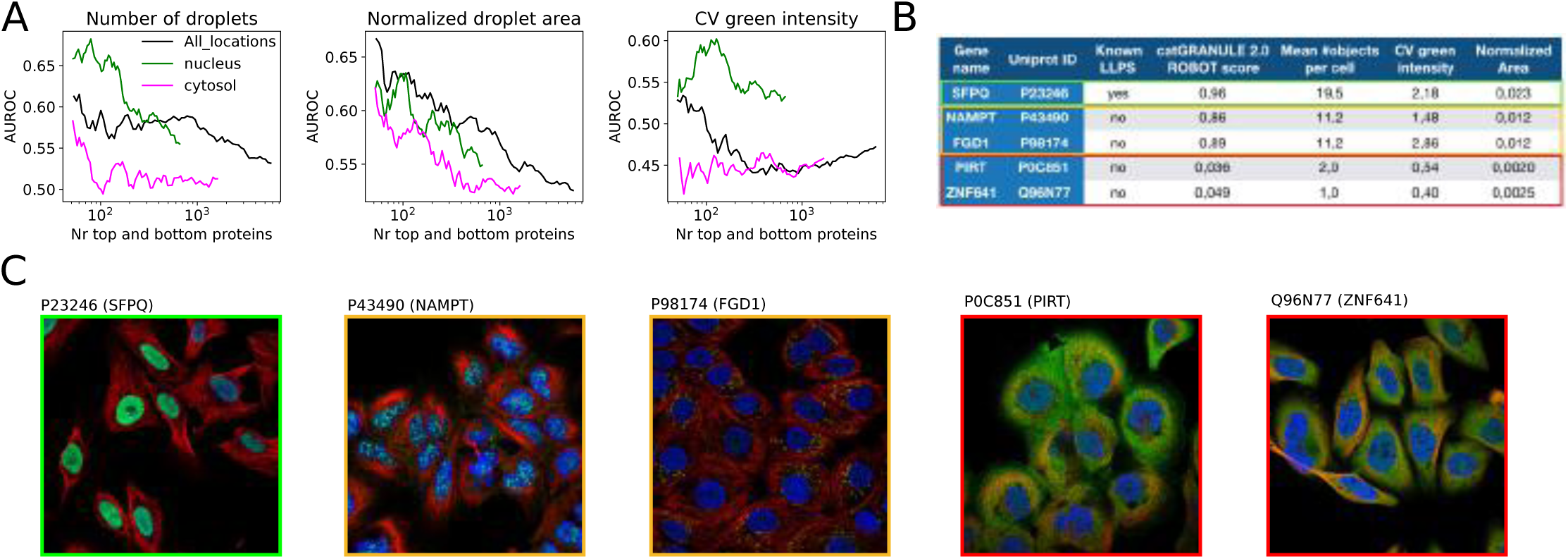
A. AUROC versus the number of top and bottom proteins, ranked according to the predicted catGRANULE 2.0 ROBOT LLPS propensity score, for the average number of droplets (i.e. green puncta, left), area of the green puncta normalized by the average area of the nuclei (center) and coefficient of variation (CV) of the green intensity over the cell (right), computed from approximately 11k antibody-based images obtained by immunofluorescence (IF) confocal microscopy from the Human Protein Atlas (HPA). Line colors indicate the selection of proteins from different subcellular locations. See the Methods section and Supplementary Table S5. B. Table showing the values of the quantities computed from the IF images for five example proteins, together with the LLPS propensity score predicted by catGRANULE 2.0 ROBOT and whether the protein was previously known to undergo LLPS. C. IF images of the proteins reported in panel B. Note that the edge color matches those in panel B.

Our predictions generally align well with the microscopy images, though not perfectly, due to the considerable variability in IF images of the same protein across different cell lines, the heterogeneity in antibody specificity, and the variability of experimental conditions.

In Figure 3B, we report the values of the features computed from the IF images shown in Figure 3C for some example proteins well known to undergo LLPS, such as SFPQ [35, 70], predicted to undergo LLPS by catGRANULE 2.0 ROBOT but not known, such as NAMPT and FGD1, and predicted to not perform LLPS (PIRT and ZNF641). We notice that the proteins predicted to undergo LLPS by catGRANULE 2.0 ROBOT show clear droplets of the green fluorescence in the nucleus (SFPQ and NAMPT) or in the cytoplasm (FGD1) (Figure 3C). Notably, FGD1 has been recently predicted between the top dosage sensitive proteins, a feature strongly associated with the ability to undergo LLPS [73]. Meanwhile NAMPT ensures the inactivation of ASK3, a protein that gets inactive after forming condensates via LLPS under hyperosmotic stress [69].

Next, we studied how the predicted LLPS score varies for proteins belonging to different subcellular locations. We found that nucleolar proteins have the highest LLPS propensity, on average, followed by cytoplasmic and nuclear proteins (Figure 4A). As expected, secreted, extracellular and membrane proteins are not predicted to undergo LLPS (Figure 4A). For the nucleolus, nucleus, cytoplasm and mitochondrion, we collected annotations for proteins from different types of liquid-like condensates from the DrLLPS database [43]. We show the distributions of the predicted LLPS score stratified by condensate in Supplementary Figure S6A. We observe that proteins belonging to the Sam68 Nuclear Body show the highest LLPS scores, on average, followed by other nuclear condensates, such as the DNA damage foci, and the nucleolus. Cytoplasmic condensates, like stress granules and P-bodies, also show high LLPS score, while the mitochondrial RNA granules are at the bottom of the ranking. These results are in agreement with recent data from filtration chromatography and dilution experiments [28]. We noticed that the composition of certain condensates largely overlaps, e.g. for stress granules and P-bodies, while others display a more unique composition (e.g. Postsynaptic density) (Supplementary Figure S6B).

**Fig. 4.**
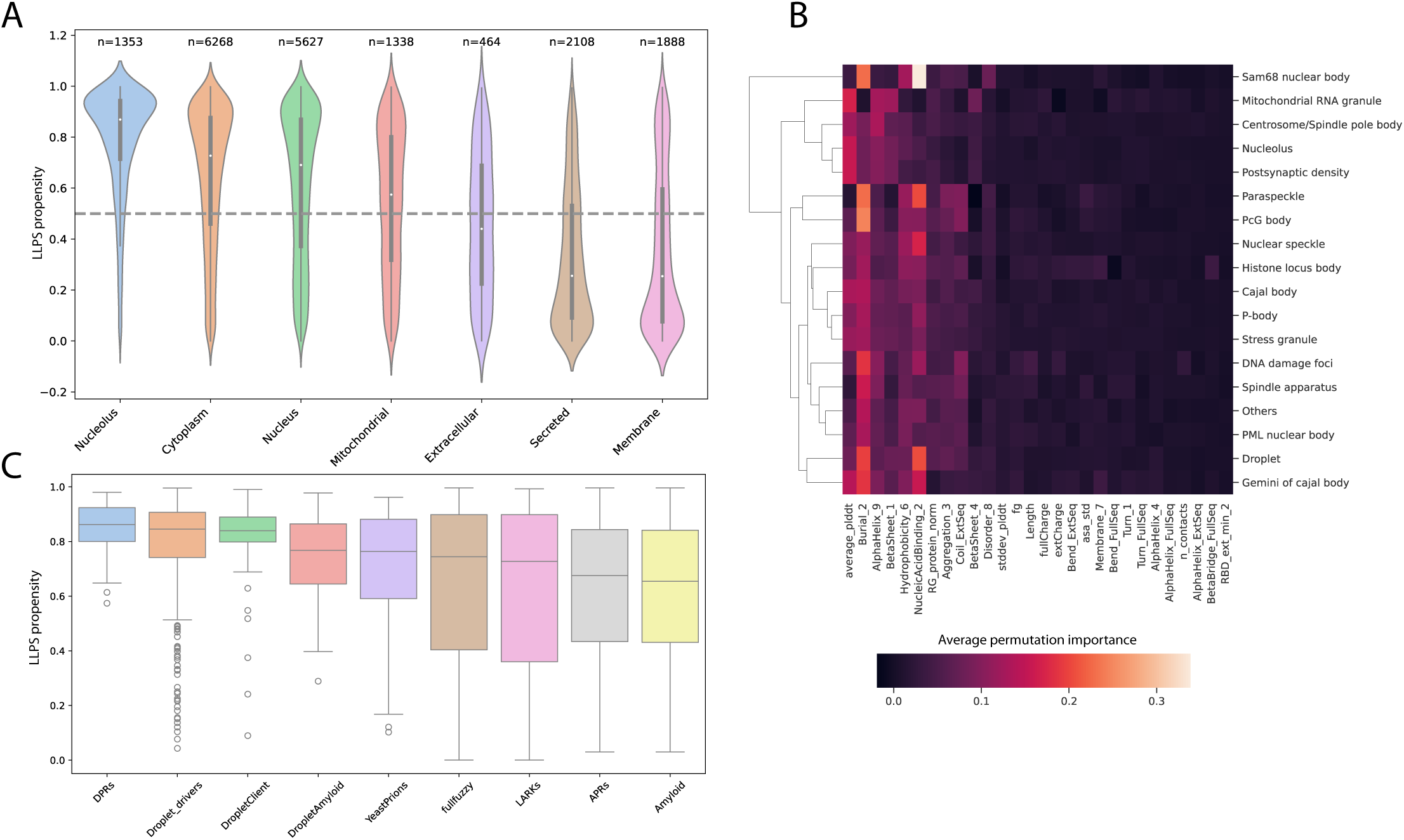
A. Violin plot showing the predicted LLPS score for proteins belonging to different subcellular locations, obtained from Uniprot [1], sorted according to descending median LLPS propensity score. The number of proteins for each subcellular location is indicated above each violin. B. Cluster map of the average permutation importance for each condensate. We show the condensates in the rows and we clustered them, we show the 28 features selected by our model ordered according to the descending permutation importance obtained from the full training dataset (see Figure 2D). C. Box plot showing the predicted LLPS propensity score for different classes of LLPS-prone proteins [66], sorted according to the median.

We then investigated the relevance of the features selected in our model for each condensate, by averaging the permutation importance over random sub-samplings of the proteins belonging to each condensate (see Methods). While for most of the condensates we found a ranking of the selected features, according to the average permutation importance, similar to the one obtained on the training dataset (Figure 2D), for the Sam68 nuclear body we observe that NucleicAcidBinding 2 is the most important feature, while structural features such as AlphaHelix 9 and Beta Sheet 1 are the top scoring for the mitochondrial RNA granule (Figure 4B and Supplementary Table S2). This pattern corresponds well with our analysis of RNA-binding proteins documented in UniProt, where 11 out of 12 proteins in the Sam68 nuclear body are RNA-binding, in contrast to 30 out of 42 in the mitochondrial RNA granule. Specifically, the Sam68 nuclear body proteins exhibited 73 instances of beta strand regions and 38 instances of compositional bias, compared to 293 beta strand occurrences and 10 instances of compositional bias in the mitochondrial RNA granule. We further confirmed the result on the RNA binding propensity by computing the catRAPID signature score for the proteins in each condensate [33], which show that proteins belonging to the Sam68 nuclear body have the highest propensity for RNA binding and there is a wide difference between condensates (Supplementary Figure S6C). Meanwhile looking at the DisProt disorder score [12] (feature Disorder 10, see Supplementary Table S2) we see less accentuated differences between condensates, although proteins belonging to the mitochondrial RNA granule have the smallest disorder score, on average (Supplementary Figure S6D).

Finally, we divided proteins in different classes according to their role in condensate formation that have been previously defined [66] (see Methods). Amyloid proteins are those found exclusively inside solid aggregates. Proteins belonging to the Amyloid-Promoting Region (APR) and Droplet-Promoting Region (DPR) classes have specific domains that can initiate amyloid or droplet formation under certain physical conditions [66]. Droplet Drivers and Clients are proteins that facilitate or participate in LLPS formation. The FullFuzzy class includes typically intrinsically disordered proteins whose interaction behaviour is dependent on the cellular context [18]. Low-complexity aromatic-rich kinked segments (LARKs) proteins are a class of proteins containing RNA binding domains [66]. Using this categorization, we first computed the predicted LLPS score for each protein and grouped the predictions by class (Figure 4C), observing a clear trend from DPRs as the highest LLPS propensity class to Amyloid as the lowest LLPS propensity class, in line with our expectations. Moreover, LARKs proteins also tend to have lower LLPS propensity scores, in line with the known role of aromatic chains in protein aggregation [15, 62]. The Drople-tAmyloid class, capable of both LLPS and LSPT, ranked intermediately, suggesting these phenomena might not be exclusive but could either compete or synergize, leading to a more thermodynamically stable structure. Indeed, while LLPS involves multivalent macromolecular interactions, LSPT also refers to specific changes in physicochemical properties. Both processes are concentration-dependent, yet intrinsic sequence features and the actual folded state of the proteins critically influence whether they undergo LLPS or LSPT [25, 67]. Additionally, we observe that some classes show a large overlap in their composition, especially the set of fullFuzzy proteins with the LARKs and DropletDrivers (Supplementary Figure S7).

These findings underscore the efficacy of our approach and its potential to advance understanding in the field of LLPS prediction and characterization.

### 2.3 LLPS profile and mutation score

We show the capability of our method to identify experimentally annotated LLPS driving regions, collected from the PhaSepDB database [75], in Figure 5. Specifically, we studied how the AUROC score varies when considering sets of top and bottom scores of increasing size (see Methods). We find that both the MLP classifier trained on structural and physico-chemical features and a Random Forest trained only on physico-chemical features achieve a AUROC *∼* 0.9 when considering the top predictions, while the performance decreases to AUROC *∼* 0.6 when taking into account all the predicted scores (Figure 5A). We noticed that the predictions of the two classifiers have a good correlation (Supplementary Figure S8A). Interestingly, we obtain better performance on proteins belonging to different organisms compared to the subset of human proteins. From this analysis we chose to adopt the Random Forest classifier, trained only on physicochemical features, as the preferred model for the computation of LLPS propensity profiles. Our choice was further supported by the superior performance, on average, of the Random Forest classifier trained on the set of physicochemical features over all the other classifiers, even when trained on the full set of features, in the prediction of the experimental LLPS-driving regions (Supplementary Figure S8B). In Figure 5B,C,E,F we show the predicted LLPS propensity profiles for four proteins from different organisms, together with the experimental LLPS driving region, finding a strong agreement between our predictions and the experimental annotation [75]. Finally, in Figure 5D we show the structure of the protein (PDB) from panel B and its LLPS propensity colored at single amino acid resolution (see Methods). In this case, the regions with higher LLPS propensity tend to match with structural elements of low complexity such as loops.

**Fig. 5.**
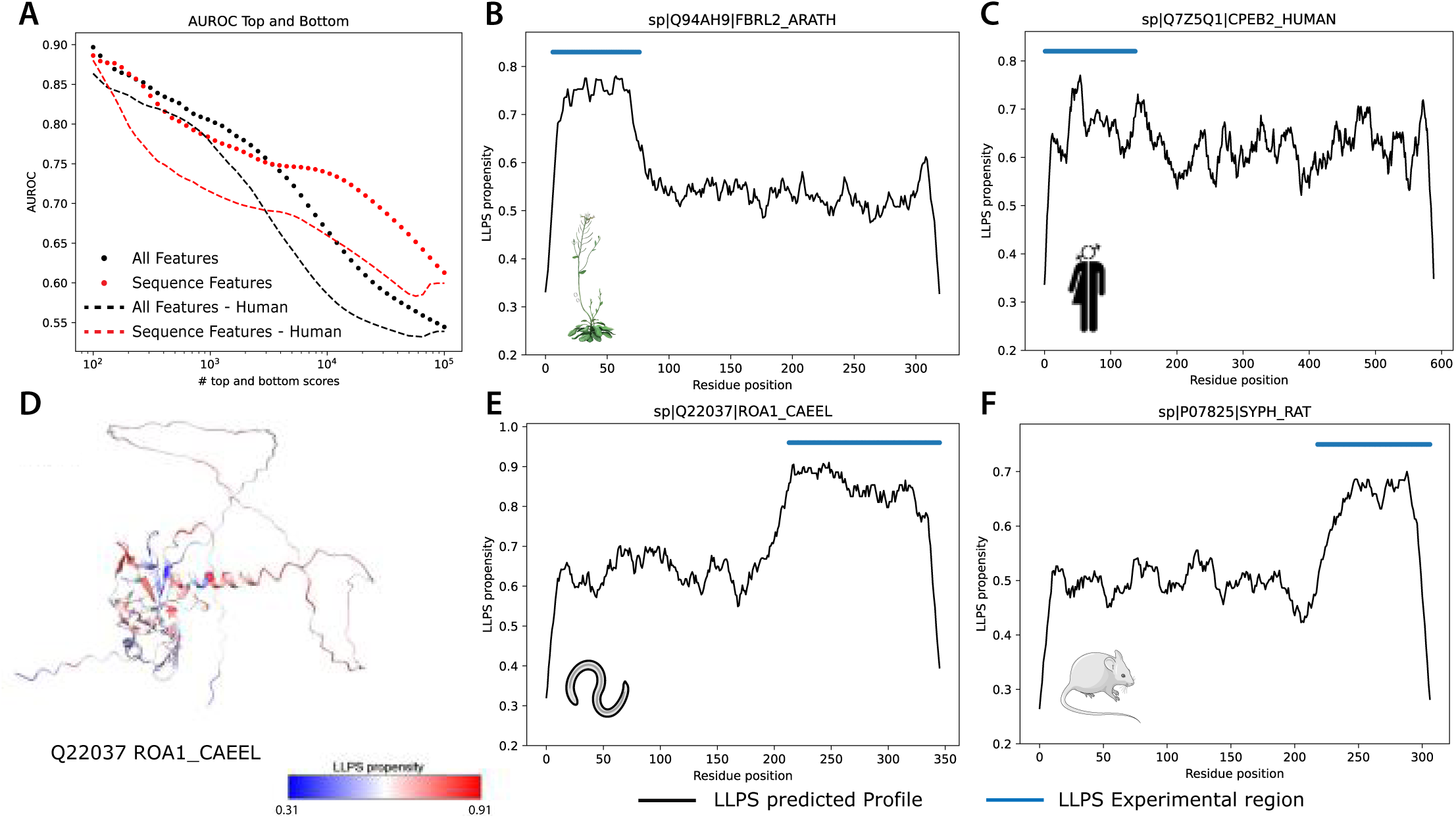
A. AUROC vs number of top and bottom scores for the MLP classifier trained on structural and physico-chemical features (black) and a Random Forest classifier trained only on physico-chemical features (red). Dots and dashes indicate proteins from all organisms or only from human, respectively. B-C-E-F. LLPS propensity profiles predicted by the Random Forest classifier trained only on physico-chemical features (black curve) and experimentally annotated LLPS driving regions (blue lines) obtained from the PhaSepDB database [75] for four proteins from different organisms. F. Protein structure colored according to the predicted LLPS propensity profile.

We exploited the capability of our method to design LLPS propensity profiles to score mutations that affect condensate formation. We highlight that the task of predicting the effect of a mutation in a protein sequence on its LLPS propensity is particularly challenging and it has not been studied extensively by previous methods. Moreover, it is known that AlphaFold2 structure prediction for single amino acid mutations is not reliable [10, 46]. For this reason, to score the effect of mutations on LLPS we use the Random Forest classifier trained only with physico-chemical features, which we have already shown that achieves comparable performance to the full model in predicting LLPS propensity profiles (Figure 5A).To score the effect of a mutation we employ the sum of the difference between the LLPS profiles of the mutant and wild-type (WT) proteins, indicated as *mut*(*i*) and *WT* (*i*), respectively, divided by the average of the profile of the WT protein

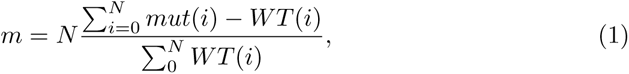

where *N* is the sequence length. Evaluating the impact of mutations on the propensity for LLPS presents significant challenges, primarily due to the dependency of experimental validation on varying environmental and cellular conditions. High-quality *in vitro* data is notably difficult to procure, as exemplified by studies such as [72, 74]. In contrast, numerous in-cell experiments, such as those reported in [22, 56], provide evidence of phase separation. These studies also include assessments ensuring that the mutations do not interfere with cellular processes or protein interactions that could sequester the protein into stress granules. Currently, the absence of a comprehensive database for extensive validation remains a major limitation in testing predictive methods. Furthermore, it is crucial to acknowledge that environmental factors like pH, ionic strength, and concentration can significantly influence the conditions under which phase separation occurs, even in mutants. Additionally, it is important to consider that cellular robustness may mitigate the impact of mutations. The presence of multiple proteins supporting the LLPS organelle and the influence of multivalency can buffer the effects of individual mutations. This suggests that in a cellular context, mutations might have a lower impact due to the collective interaction of various molecules involved in the LLPS process.

To validate the capability of our method in identifying the effect of mutations on LLPS, we collected a list of 24 distinct mutations of 9 proteins that undergo LLPS in the WT form but show increased or reduced LLPS propensity when mutated (see Methods and Supplementary Table S6), and we compare the performance in scoring mutations of catGRANULE 2.0 ROBOT with those achieved by catGRANULE 1.0 [7] and PSPHunter [59].

First, we compute the LLPS score of the WT proteins and we notice that only catGRANULE 2.0 ROBOT correctly predicts all of them to undergo LLPS (Figure 6A). Next, we compute a mutation score for the 24 mutations for the three algorithms and we evaluate the fraction of correctly predicted mutations separately for the set of mutations increasing or decreasing the LLPS propensity. We found that catGRANULE 2.0 ROBOT and catGRANULE 1.0 correctly predict the 80% of the mutations with a negative effect on LLPS, while PSPHunter correctly calculates only the 50%. For mutations with a positive effect on LLPS catGRANULE 2.0 ROBOT outperforms the other algorithms (Figure 6B).

**Fig. 6.**
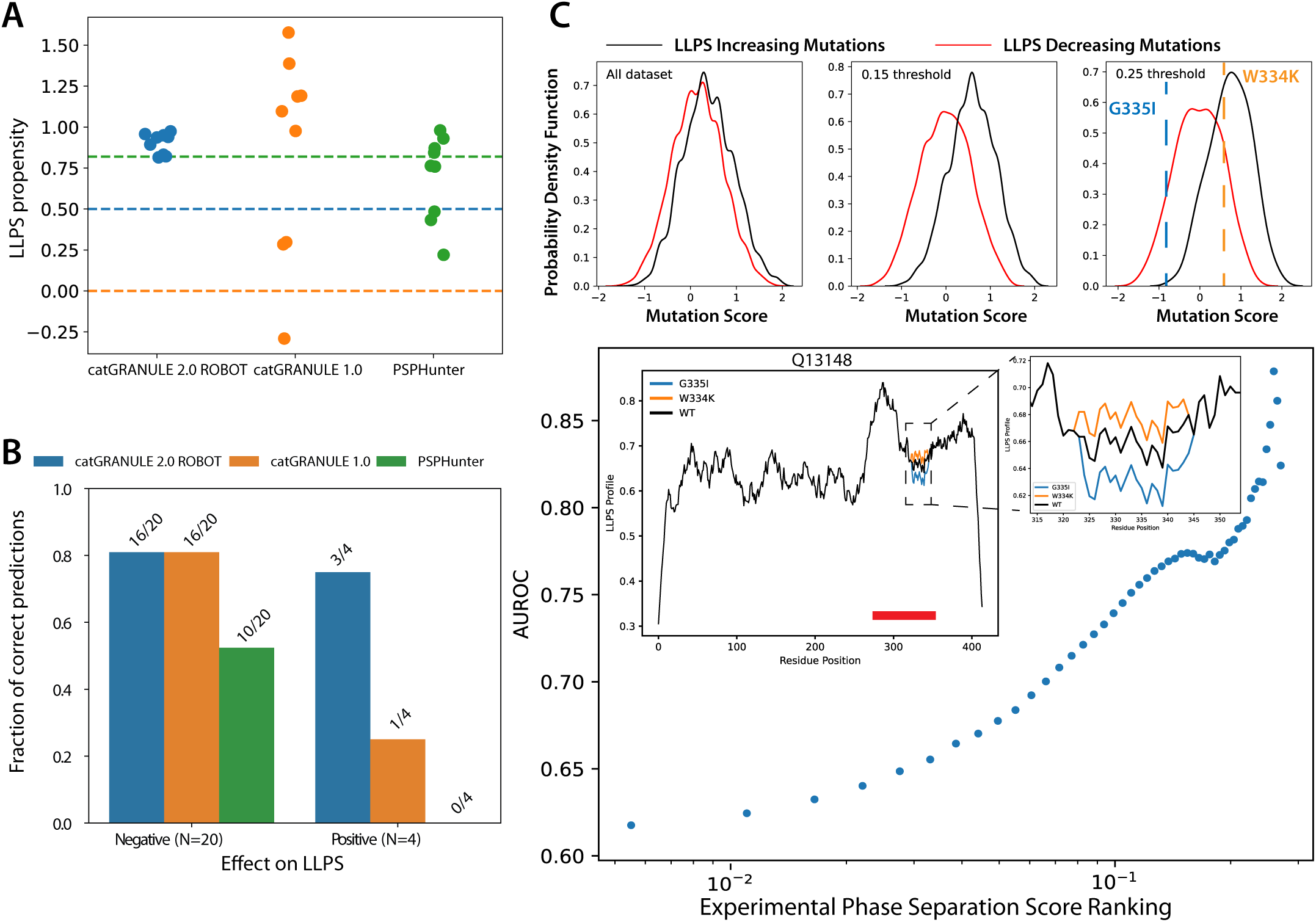
A. LLPS propensity score computed by catGRANULE 2.0 ROBOT, catGRANULE 1.0 and PSPHunter on the WT sequence of 9 proteins for which mutations affecting LLPS were collected. Colored dashed lines indicate the threshold to discriminate LLPS from non-LLPS proteins, with the color matching the algorithm. B. Bar plot showing the fraction of correctly predicted mutation scores by catGRANULE 2.0 ROBOT, catGRANULE 1.0 and PSPHunter, for a set of 24 mutations including 20 mutations with a negative effect on LLPS and 4 mutations with a positive effect. C. Distributions of the catGRANULE 2.0 ROBOT mutation score for mutations decreasing or increasing LLPS (red and black curves, respectively) from a mutational scanning of TDP-43 [8], at different thresholds on the experimental phase separation score. In the rightmost panel the colored dashed lines show the predicted mutation score of two selected mutations. D. AUROC computed on the catGRANULE 2.0 ROBOT mutation scores for the mutational scanning of TDP-43, as a function of the threshold on the experimental phase separation score. Increasing the threshold corresponds to selecting more restricted sets of mutations, with stronger positive and negative experimental effect on LLPS. The inset shows the LLPS propensity profiles predicted by catGRANULE 2.0 ROBOT for the WT sequence of TDP-43 and the two mutations shown in panel C. The red line indicates the experimental LLPS region.

Finally, to assess the ability of catGRANULE 2.0 ROBOT to predict the effect of mutations on LLPS propensity under consistent environmental conditions, we analyzed a mutational scanning dataset of TDP-43 [8], a protein known to undergo LLPS and that is implicated in neurodegenerative diseases. This large-scale screening highlights an astonishing correlation between the formation of LLPS and cellular toxicity. Specifically, mutations that facilitate LLPS in TDP-43 increase cellular toxicity due to interactions with other cellular molecules. Conversely, mutations leading to LSPT result in less toxic, more inert protein aggregates [8]. The mutational scanning includes approximately 60000 mutations of TDP-43, including both single and double mutations [8] (see Methods). In Figure 6C we show the distribution of the mutation score (Eq. (1)) predicted by catGRANULE 2.0 ROBOT, considering the full mutational scanning or restricting the absolute value of the experimental phase separation score to a certain threshold. We observe that the separation between the distributions of LLPS decreasing and LLPS increasing mutations, represented by the red and black curves, respectively, increases with the threshold on the experimental score, and that catGRANULE 2.0 ROBOT correctly predicts the sign of the mutation score for a LLPS decreasing (G335I) and a LLPS increasing (W334K) mutations whose physical properties were studied in detail [8] (Figure 6C). In Figure 6D we show the AUROC achieved by catGRANULE 2.0 ROBOT, computed on sets of top and bottom mutations selected at different thresholds of the experimental phase separation score. While at low values of the threshold catGRANULE 2.0 ROBOT achieves an AUROC score slightly higher than 0.6, increasing the threshold, which corresponds to selecting sets of mutations with stronger negative or positive effect on TDP-43 LLPS, it reaches AUROC close to 0.9 (Figure 6D). The inset shows the profiles of the TDP-43 WT protein, which nicely agrees with the experimental LLPS region (red line) [24], and of the two mutations already shown in Figure 6C. Furthermore, we observed that the histograms of the catGRANULE 2.0 ROBOT mutation score for single and double mutations are both approximately Gaussian centered at zero (Figure S8C). Nevertheless, for double mutations the tails of the distribution are heavier, suggesting that they can have a stronger effect on the LLPS propensity compared to single mutations, as expected [8].

Overall, we showed that catGRANULE 2.0 ROBOT can accurately predict LLPS propensity profiles and LLPS-prone regions of proteins from different species. Moreover, it outperforms previous methods in scoring the effect of single and multiple amino acids mutations on LLPS, making it an appealing tool for the design of proteins or peptides with tunable LLPS properties.

## 3 Discussion

The emergence of LLPS as a central molecular process governing membrane-less organelle formation within cellular environments underscores its profound implications in human health, particularly in the context of neurodegenerative disorders. Unlike the irreversible aggregation characteristic of LSPT, LLPS condensates present a reversible but less thermodynamically stable state, facilitating dynamic protein and RNA compartmentalization and influencing cellular activities [79].

Here, we have developed an advanced LLPS protein predictor, catGRANULE 2.0 ROBOT. This tool integrates structural and sequence-based features from AlphaFold2 models, enabling detailed profiling of phase separation properties and mutant computations. Notably, the predictor includes a user-friendly web interface and pre-calculated scores for various model systems. A number of improvements position catGRANULE 2.0 ROBOT at the forefront of predictive tools for studying protein phase separation, with capabilities that surpass those of current methodologies by integrating comprehensive biophysical data and cutting-edge computational predictions. Our integrated approach is expected to refine our understanding of LLPS processes and facilitate the exploration of novel therapeutic and biological insights [79].

In detail, a notable aspect of our study is the capability to identify and predict the effects of mutations on LLPS propensity. Through a comprehensive analysis, we demonstrated the efficacy of our predictor in accurately spotting single mutations and their impact on phase separation tendencies. This ability holds great promise for elucidating the molecular mechanisms underlying disease-associated mutations and guiding precision protein engineering efforts [59, 79]. Most importantly, our approach enables the generation of detailed LLPS propensity profiles along protein sequences at single amino acid resolution. The ability to identify phase separation-driving regions enhances our understanding of protein structure-function relationships and provides valuable guidance for experimental design and interpretation [21].

Through a combination of direct and indirect experimental approaches [8, 63], we have corroborated the predictions, affirming the robustness and reliability of catGRANULE 2.0 ROBOT. The analysis of deep mutational scanning, particularly in the case of TDP-43 [8], further strengthens the alignment between our predictions and experimental observations, underscoring the clinical relevance of our study.

catGRANULE 2.0 ROBOT exploits AlphaFold2 models to integrate structural and sequence-based features, enabling detailed profiling of phase-separating proteins and computation of mutants. A key enhancement is the use of AlphaFold2 [27], with plans to upgrade to AlphaFold3 [2] in future iterations for even more precise structural predictions. Additionally, we aim to integrate it with algorithms such as catRAPID [4] and scRAPID [16] to better predict specific RNAs that contribute to protein crowding, enhancing the model’s ability to simulate complex biological environments, and the cell type specific expression of the RNAs interacting with proteins undergoing LLPS. We also plan to include context-dependent features such as ions and cellular-specific chemical modifications, which greatly impact phase separation through mechanisms like ionic strength, pH, and concentration. Further experimental validation through techniques such as FRAP and FCS [49] will be essential to link LLPS-related properties with specific structural features now predictable with AlphaFold3 [2]. These developments will position the new generation of algorithms at the cutting edge of predictive tools for studying protein phase separation, offering unparalleled accuracy and comprehensive biophysical data integration.

## 4 Conclusion

Here we predicted the phase separation propensities for a number of proteins residing in various cellular compartments, including stress granules, nuclear bodies, Cajal bodies, and P-bodies. The physico-chemical conditions under which these proteins are expressed, such as environmental factors and protein concentration, are crucial for the formation of different types of phase-separated organelles. We anticipate that further investigation into these conditions, along with chemical modifications of proteins and RNAs, may lead to the discovery of other types of organelles. For this reason, we plan to enhance our predictions by integrating data on protein-protein and protein-RNA interactions. This aims to provide a clearer understanding of the compositions of various types of assemblies. Our approach is somewhat reminiscent of previous efforts to identify proteins interacting with RNA-binding proteins [9]. In a recent endeavor in this direction [14], we have revealed important characteristics of phase separation. By continuing to explore the complex interplay between protein concentration, environmental conditions, and protein interactions, we hope to uncover new mechanisms and principles underlying cellular organization and function [79]. In conclusion, our study not only advances our understanding of LLPS but also holds significant implications for protein engineering. The ability to precisely manipulate LLPS propensity at the molecular level opens avenues for the design of novel proteins and mutations with tailored phase separation behavior, offering unprecedented opportunities for therapeutic innovation [57, 78]. To facilitate the use of catGRANULE 2.0 ROBOT by others, we have developed a web server, available at https://tools.tartaglialab.com/catgranule2. This tool allows the community to explore LLPS predictions and design mutants, offering significant potential for protein engineering and therapeutic innovation.

## 5 Methods

### 5.1 Construction of the training dataset

To build the positive set of LLPS-prone proteins, we collected data for human proteins from several databases and resources. Specifically, we used:

- 929 proteins annotated as “Droplet state” in the PRALINE database [65];
- 3876 proteins from the CD-CODE database [53];
- 3807 proteins used in a recent study on LLPS predictors (provided in table S1 from [31]);
- 117 proteins defined in [76] (obtained from Table S3 in [30], where they are defined as the “Gingras gold standard”);
- 3633 proteins from the DrLLPS database [43];
- 833 proteins from the PhaSepDB database (v2.1) [75];
- 59 proteins from the PhaSePro database [39];
- 92 proteins from LLPSDB [68].

We considered the union of these sets since some of them have a large overlap, as shown by the Venn diagram in 1B, obtaining 5656 proteins. Next, we used CD-HIT [17] to filter proteins from the positive set for 50% sequence identity, obtaining 4807 proteins. To generate the negative set, we removed the proteins belonging to the positive set from the human proteome, and we also removed their first interactors based on protein-protein interactions collected from the BioGRID database (v3.5.175) [45]. Finally, we filtered proteins in the negative set using CD-HIT, as described above. To ease the comparison with the top performing state-of-the-art methods, which are MaGS [30, 31], PICNIC and PICNIC-GO [23], we included in our training positive set the LLPS-prone proteins on which those algorithms were trained. Since the positive training set for PICNIC was extracted from the CD-CODE database [53], but the authors do not provide the proteins that have been used in the training, we included in the positive training set all the proteins belonging to CD-CODE, after the CD-HIT filtering described above. Next, we randomly sampled the same amount of negative proteins. Considering the intersection with the proteins for which a pdb file is present in the AlphaFold Protein Structure Database (https://alphafold.ebi.ac.uk/) [64], the final training dataset is made of 3333 positive and 3252 negative proteins. We used the remaining LLPS-prone proteins as an independent positive test set and we sampled the negatives as for the training dataset. We obtained a test dataset with 1422 positive and 1376 negative proteins.

We used the Panther database to represent the composition of the training dataset in terms of Biological Process (BP), Molecular Function (MF) and Protein Class (PC) Gene Ontology (GO) categories [40]. We used a chi-square test to quantify the enrichment in the positive or negative training set versus the rest of the proteome, or in the positive versus the negative set, for the groups of categories mentioned above.

We also conducted a functional enrichment analysis on the positive and negative proteins of the training dataset using gProfiler [51]; the results are reported in Supplementary Table S1.

Next, from the training dataset we computed the correlation matrix of the features and we represented it in a hierarchical cluster map using the function “clustermap” of the “seaborn” Python package.

### 5.2 Definition of model features

#### 5.2.1 Physico-chemical features

We use a list of 80 experimental scales encoding physico-chemical properties of proteins that describe aggregation, hydrophobicity, membrane, nucleic acid binding, disorder, burial, alpha helix, beta sheet, turn propensities, and two phenomenological sequence patterns [7, 29]. These features are computed solely based on the protein sequence and they were already used in the cleverSuite [29] and in the training of the catGRANULE 1.0 algorithm for LLPS propensity prediction [7]. Each protein sequence is transformed in a list of 82 numbers, representing the average of each physico-chemical feature over the sequence.

#### 5.2.2 AlphaFold2-derived features

We downloaded the structures predicted by AlphaFold2 [27] for the whole human proteome from the AlphaFold Protein Structure Database (https://alphafold.ebi.ac.uk/) [64]. For the proteins having multiple predicted models we select the model with highest average predicted local-distance difference test (pLDDT), a measure of local accuracy of the AlphaFold2 prediction. Next, we used the ”Bio.PDB” submodule of the ”Biopython” (version 1.81) python package to extract structural features from the pdb file for each protein. Specifically, we extract the number of contacts, the radius of gyration (RG) and the accessible surface area (asa), from which we define the exposed residues of the protein as those with *asa >* 0.5, while the rest are considered as internal. Next, for the internal and the exposed residues we extract the secondary structure properties (helix, strand, coil, turn, accessibility, disorder) and the charge.

We also include features encoding the RNA-binding propensity of the residues in the internal, exposed or full protein by leveraging a set of experimentally determined RNA binding domains (RBDs) [13]. We apply a sliding window on each protein sequence taking into account its exposed and internal parts, and we compute the distance in composition to each experimental RBD. We collect as features the distances and the indices of the 3 RBDs with smallest distances. Considering the internal and the exposed parts of the protein as well as the whole sequence, we obtain 18 features in total.

Finally, we include the average and the standard deviation of the pLDDT as additional features, as well as the number of contacts and radius of gyration normalized by the protein length. For each protein structure we obtain a vector of structural and RNA-binding features of length 46; adding the 82 physico-chemical features we have 128 features per protein.

In Supplementary Table S2 we report a mapping of the feature names and their categorization in feature families. We also report the original references from which the physico-chemical scales were obtained.

### 5.3 Model training

We use the Python library scikits-learn (version 1.1.1) [48] to train multiple classifiers for LLPS propensity prediction. Specifically, we use Decision Tree, Random Forest, XGBoost, Gradient Boosting, Gaussian Naive Bayes, TabNet, Logistic Regression, catBoost, K-neighbors and a multilayer perceptron (MLP). For each classifier we define a scikits-learn pipeline including feature scaling (“StandardScaler” function), feature selection and training of the classifier. We use ElasticNet as the feature selection method since it performs multivariate feature selection, i.e. it takes into account the (possibly non-linear) statistical dependence between features when selecting them during training [81]. To avoid overfitting, we use a 5-fold cross validation for training (“gridsearchCV” function), including a grid search for hyperparameter optimization on both the feature selection method and the classifiers. To this end, we defined a parameter search space specific to each classifier. We trained each classifier using the Area Under the Receiver Operating Characteristic Curve (AUROC) as the scoring function. We selected the MLP as the best model based on the AUROC obtained on the independent test set. In the feature selection step, we obtained a set of 28 features selected by ElasticNet. We computed the Spearman’s correlation coefficient between each feature and the predicted LLPS propensity score on our training dataset, to obtain a sign for the effect of a feature increase or decrease on the LLPS propensity score. To quantify the contribution of each selected feature to the final AUROC score, we used the “permutation importance” function from the “sklearn.inspection” submodule, which randomly permutes a feature in the dataset and computes the AUROC using the trained model. We represent the distributions of the values of the permutation importance obtained from 50 repetitions of the described procedure using a box plot. For the 10 features with highest permutation importance, we performed a Wilcoxon’s rank-sum test (python function “ranksums” from the “scipy.stats” submodule) between the positive and negative sets of proteins belonging to the training dataset.

To test the performance of the feature selection done using ElasticNet, we iteratively selected the top *k* features using a univariate method (”SelectKBest” from scikits-learn), with *k* ranging from 1 to the total number of features in the dataset (*N* = 128), and then trained a MLP classifier. Moreover, we also trained a linear model using all the features.

Finally, we recursively eliminated the features selected by ElasticNet from the training dataset and we trained a MLP classifier each time. We represented the selected features at each iteration in terms of the feature families defined in Supplementary Table S2 using a stacked area chart.

### 5.4 Model validation

We compared the performance of catGRANULE 2.0 ROBOT on the independent test set with those of catGRANULE 1.0 [7] and of the four top performing state-of-the-art algorithms, MaGS [30, 31], PICNIC, PICNIC-GO [23] and PSPHunter [59].

#### 5.4.1 catGRANULE 1.0

catGRANULE 1.0 is a sequence-based predictor of LLPS propensity [7]. It uses the RNA-binding propensity, structural disorder content, the frequencies of arginine, glycine, phenylalanine and sequence length as features. It defines a granule propensity of the amino acid at each position in the protein sequence as a linear combination of the features, computed over windows of length 7. The coefficients of the linear combination were optimized via a Monte Carlo method. The algorithm was trained on a dataset of yeast proteins.

#### 5.4.2 MaGS

MaGS (Membraneless organelle and Granule Score) uses sequence-based protein features and others, such as protein abundance, number of annotated interaction partners, number of annotated phosphorylation sites etc) to encode each protein into a feature vector [30, 31]. The algorithm is based on a generalized linear model trained over a curated set of proteins obtained from the literature.

#### 5.4.3 PICNIC

PICNIC (Proteins Involved in CoNdensates In Cells) is a machine learning-based model, based on a catBoost classifier, that predicts the LLPS propensity of proteins [23]. It uses both sequence-based (including sequence complexity, IUPred disorder score and amino acid co-occurrences) and structural features derived from AlphaFold2 models. PICNIC-GO includes an extended set of features based on Gene Ontology terms.

#### 5.4.4 PSPHunter

PSPHunter (Phase-Separating Protein Hunter) is a machine-learning algorithm that predicts protein LLPS propensity and identifies the key residues for LLPS [59]. It integrates sequence-based and functional features of the protein, including amino acid composition, evolutionary conservation, functional site prediction, protein network properties retrieved from protein-protein interaction networks, and word embedding vectors obtained using the word2vec method. PSPhunter is trained on several training datasets of small size using proteins from different species.

We computed the ROC curves using the function ”roc curve” from ”sklearn.metrics” and the AUROC using the function ”roc auc score” on the test dataset.

For the computation of the catGRANULE 2.0 ROBOT LLPS propensity score in other species, we retrieved proteins annotated as LLPS-prone from the DrLLPS database [43], we downloaded the pdb files for those proteins from the AlphaFold Structure Database [64], we encoded each protein into the vector of 128 features described above and we predicted the LLPS propensity score using our pre-trained MLP classifier. We retrieved the numbers of correctly predicted proteins by PICNIC from [23] and, for the species that are predicted both in our study and by PICNIC, we tested, for each species separately, if the fraction of correctly predicted LLPS-prone proteins by catGRANULE 2.0 ROBOT was significantly larger than the one predicted by PICNIC using a Fisher’s exact test (function ”fisher exact” in scipy.stats). Regarding the analysis of the LLPS propensity per condensate, first we retrieved the annotation of the protein sub-cellular locations from Uniprot (https://www.uniprot.org/) [1] and the annotation of LLPS condensates from the DrLLPS database [43]. Since proteins can be found in multiple condensates, we represented the intersections using an upset plot (”upset” function from the ”UpsetR” package, version 1.4.0, in R version 4.0.3). The violin plots are obtained using the function ”violinplot” from the seaborn Python package.

To test the importance of the features selected by ElasticNet for each condensate, we repeated the computation of the permutation importance previously described for the sets of proteins belonging to each condensate. Since the permutation importance measures the contribution of each feature to the chosen scoring function, which is the AUROC in our case, for each condensate we generated a negative set of the same size of the positive set by randomly sampling proteins from our full negative set. We repeated this procedure 50 times for each condensate and we represented the average permutation importance for each condensate and each of the 28 features selected by ElasticNet using the ”clustermap” function in seaborn. We clustered the rows to show the similarity between condensates, while we ordered the columns (features) according to the values of the permutation importance obtained from the training dataset. Finally, we show the RNA binding propensity, quantified by the catRAPID signature score [33], and the DisProt disorder score (feature Disorder 10 according to our nomenclature, see Supplementary Table S2) [12] of proteins per condensate in two boxplots, computed using the function ”boxplot” in seaborn.

#### 5.4.5 Validation on Immunofluorescence images from the Human Protein Atlas

For the analysis of antibody-based images obtained by immunofluorescence (IF) confocal microscopy from the Human Protein Atlas (https://www.proteinatlas.org/ humanproteome/cell) [63], we retrieved a curated list of 11608 images from [77]. We used a CellProfiler3 [38] pipeline provided in [77], which we adapted to compute additional quantities from the images and to use it with CellProfiler4.2.6 [58].

Specifically, we perform cell segmentation from the IF image through the Otsu’s thresholding method using the red and blue channels, which quantify the microtubules and the DAPI, respectively. Then, we compute the standard deviation and the mean of the green intensity per cell, whose ratio defines the coefficient of variation (CV). Next, we segment the nuclei using the blue channel, we compute the area of each nucleus and we take the average over each image. Finally, we segment the droplets (i.e., puncta of the green fluorescent protein) for each cell using the robust background thresholding method, and we compute the area, measured in pixels, of each droplet, which is made adimensional by dividing it by the average area of the nuclei for the corresponding image. To obtain measurements at the protein level and compare them to the LLPS propensity scores predicted by catGRANULE 2.0 ROBOT, we computed, for each protein, the average number of droplets, the maximum normalized area and the maximum CV of the green signal. In this way we obtained the measurements for 10757 IF proteins, each corresponding to one IF image. We provide all the computed scores in Supplementary Table S5.

### 5.5 Computation of the LLPS propensity profiles and prediction of the effect of mutations on LLPS

catGRANULE 2.0 ROBOT predicts the LLPS propensity of a protein based on a set of sequence- and structural-based features. However, to compute a LLPS propensity profile and to allow the usage of our model on deep mutational scanning of proteins, which generate tens of thousands of mutations, we chose to train the model using only physico-chemical features. This choice allows a fast analysis of deep mutational scanning of proteins and it is supported by previous studies that showed that AlphaFold2 cannot predict reliably the structure of proteins subjected to single-point mutations [10, 46].

#### 5.5.1 Prediction and validation of LLPS propensity profiles

We generate a LLPS propensity profile for a protein by applying a sliding window to a protein sequence and scoring each segment with the trained model. In this way we obtain a LLPS propensity score at single amino acid resolution. We trained different classifiers using both the full set of features or only the set of physico-chemical features. Then, we collected approximately 250 proteins from different organisms from the PhaSepDB database [75], where regions responsible for LLPS are annotated over the sequence. Furthermore, to increase the accuracy and the sensitivity of the prediction, we filtered out proteins with more than 90% of the sequence annotated as the LLPS-prone region from the PhaSepDB dataset. Using this dataset, we found that the optimal size of the sliding window for the computation of the LLPS propensity profiles is 21 amino acids. To quantify the performance of the trained models over the PhaSepDB dataset, we concatenated all the protein sequences and we ranked the aminoacids according to the predicted LLPS propensity. Next, we selected subsets of top and bottom LLPS propensity scores and we computed the AUROC score, where the true classes are obtained from the PhaSepDB database (see Figure 5A). To select a model for the computation of the profiles, we compared the average AUROC obtained from the top-bottom scores approach and we found that a Random Forest classifier, trained only on the set of physico-chemical features, achieves the best performance (Supplementary Figure S8B). Moreover, the average of the LLPS propensity profiles obtained with the Random Forest classifier shows a good correlation, quantified by the Spearman’s correlation coefficient, with the LLPS propensity score predicted by the full model, which is the MLP classifier trained on the full set of features (Supplementary Figure S8A).

#### 5.5.2 Prediction of the effect of mutations on LLPS propensity

We collected a list of 24 mutations of 9 proteins, including single and multiple amino acid mutations, from the literature and the Uniprot database [1]. They are reported in Supplementary Table S6, where we also indicate the references from which the mutations have been retrieved. These mutations were categorized according to their annotated effect of increasing or decreasing LLPS propensity and - or affecting the protein localization in SGs and PBs. Next, we predicted the LLPS propensity profiles of the wild-type (WT) and mutated proteins, and we computed a mutation score as defined in Eq (1). We also computed the LLPS propensity of the WT and mutated proteins using the PSPHunter [59] and catGRANULE 1.0 [7] web servers, and we compared the fraction of mutations for which the effect on LLPS propensity was correctly predicted, separately for mutations decreasing and increasing the LLPS propensity, between catGRANULE 2.0 ROBOT and these two algorithms (Figure 6B). The predicted scores for the WT and mutated proteins for the three algorithms are reported in Supplementary Table S6.

Finally, we considered a mutational scanning of TDP-43 where approximately 60000 mutations of the prion-like domain were generated and their toxicity was quantified in yeast cells [8]. The authors showed that mutations that increase protein aggregation strongly decrease toxicity, while toxic mutations promote LLPS. Thus, the toxicity score can be used as a proxy of experimental LLPS propensity. We predicted a LLPS propensity profile for each mutation and we computed a mutation score as described above. Then, we employed a top-bottom approach as we did for the validation of the LLPS propensity profiles. Specifically, we ranked the mutations according to the experimental phase separation score and we set different thresholds on this score. For each threshold we computed the AUROC, as shown in Figure 6C.

## Supplementary information

Additional File 1. Supplementary Figures S1-S8.

Table S1. Results of a functional enrichment analysis on the positive and negative proteins belonging to the training dataset using gProfiler.

Table S2. Mapping of feature names and categorization in feature families.

Table S3. LLPS scores predicted by catGRANULE 2.0 ROBOT for the whole human proteome, with annotation of proteins belonging to the training and test datasets and if the proteins were already known to undergo LLPS.

Table S4. LLPS scores predicted by catGRANULE 2.0 ROBOT for the sets of proteins annotated as LLPS-prone in the DrLLPS database for different species.

Table S5. Features computed from immunofluorescence microscopy images obtained from the Human Protein Atlas, aggregated at the protein level.

Table S6. List of 24 mutations affecting LLPS curated from the literature with predicted mutation and WT scores by catGRANULE 2.0 ROBOT, catGRANULE 1.0 and PSPHunter.

## Declarations

### Ethics approval and consent to participate

Not applicable.

### Consent for publication

Not applicable.

### Availability of data and materials

All the data needed to reproduce the analysis in this manuscript are available from the supplementary material. The catGRANULE 2.0 ROBOT web server is available at https://tools.tartaglialab.com/catgranule2.

### Authors’ contributions

MM, JF and GGT conceived the study. MM and JF developed the method and performed the computational analyses. DMV and GB contributed to the computational validation of the method. AA developed the catGRANULE 2.0 ROBOT web server. MM, JF and GGT wrote the manuscript. TC and NSdG provided critical insights and edited the manuscript. GGT supervised the study.

## Acknowledgements

The authors acknowledge Andrea Vandelli and Jakob Rupert for useful discussions.

