## Supplementary Materials and Figures for "Accurate Predictions of Liquid-Liquid Phase Separating Proteins at Single Amino Acid Resolution"

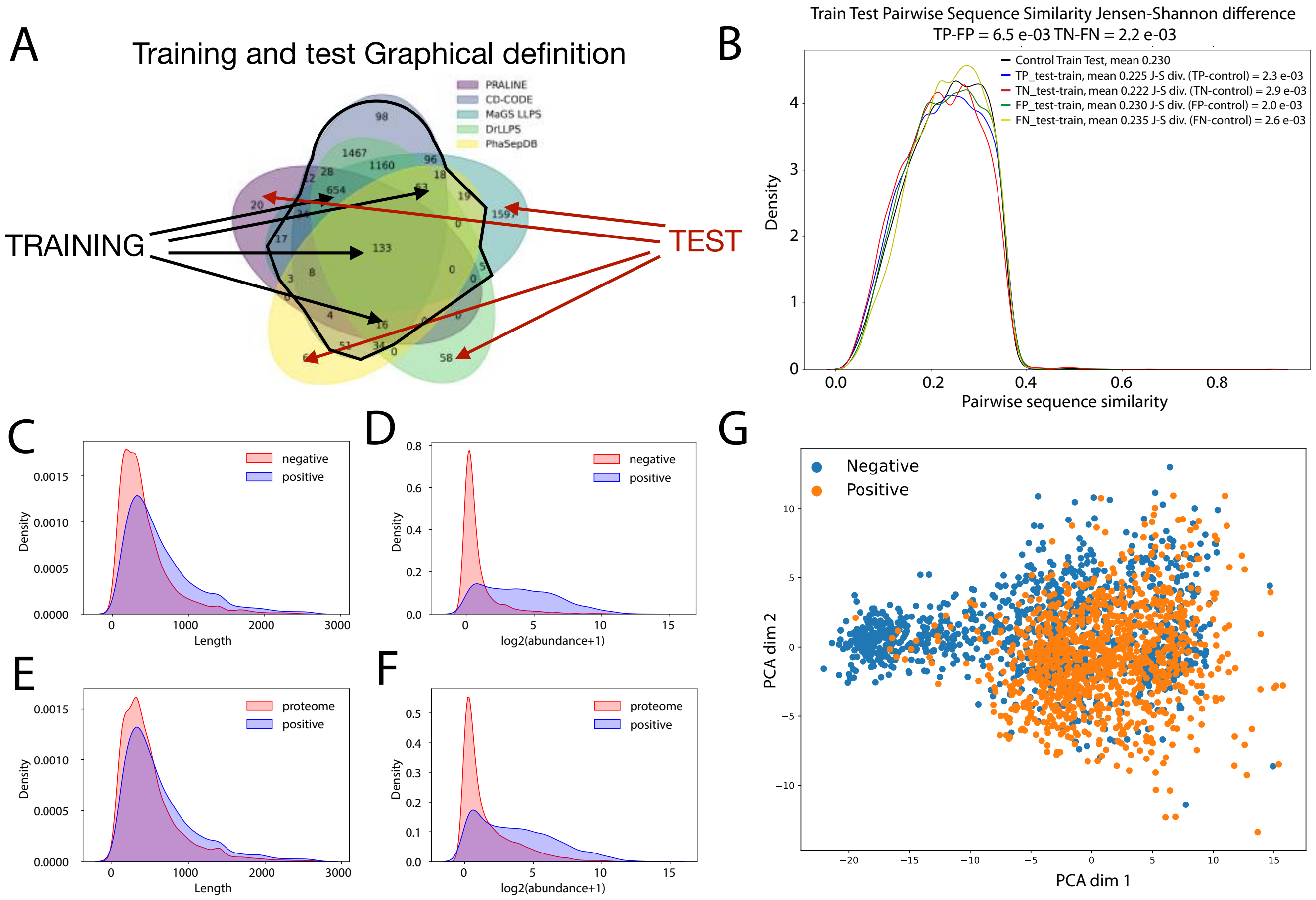


FIG. S1: A. Venn diagram showing the overlap of LLPS-prone proteins collected from different databases with highlighted selection of the positive set of proteins for the training and test datasets. B.The distribution of pairwise sequence similarity was computed for both the training and test datasets, including true positives, false positives, true negatives, and false negatives. The results show that the distributions of pairwise alignment are similar across these categories. The values of the Jensen-Shannon distance between the distribution of true and false positives with the true and false negatives are shown in the legend, highlighting the absence of sequence composition biases in catGRANULE 2.0 ROBOT predictions. C. Probability density of protein expression (obtained from the PaxDB database `https://www.pax-db.org/` [1]) for LLPS-prone proteins (blue) and the negative training set (red). D. Probability density of protein length for LLPS-prone proteins (blue) and the negative training set (red). E-F. Same as panels C-D for LLPS-prone proteins (blue) and the rest of the proteome (red). G. Principal Component Analysis (PCA) plot of the positive and negative proteins computed using the set of 128 features used by catGRANULE 2.0 ROBOT to encode proteins, highlighting the presence of a partial separation between the positive and negative sets along PC1.

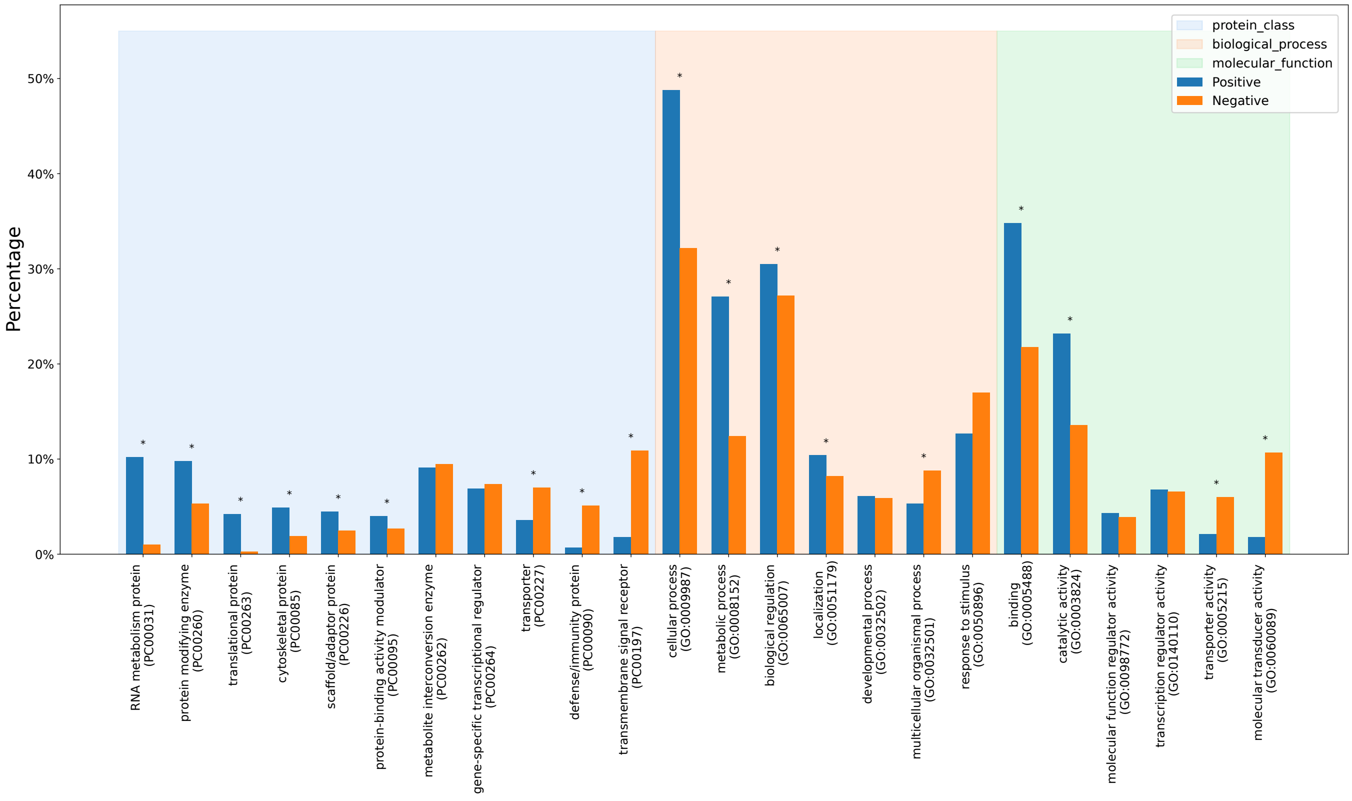


FIG. S2: Bar plot showing the composition of the catGRANULE 2.0 ROBOT training dataset in terms of categories obtained from the Panther database. The coloured regions highlight different classifications of the proteins, namely Protein Class (light blue), Biological Process (light orange) and Molecular Function (light green). Blue and orange bars represent the percentage of proteins from the positive and negative set, respectively, belonging to a specific Panther class. The stars indicate a p-value smaller than 0.05 from a chi-square test between the fraction of proteins in the positive vs negative sets for a given category.

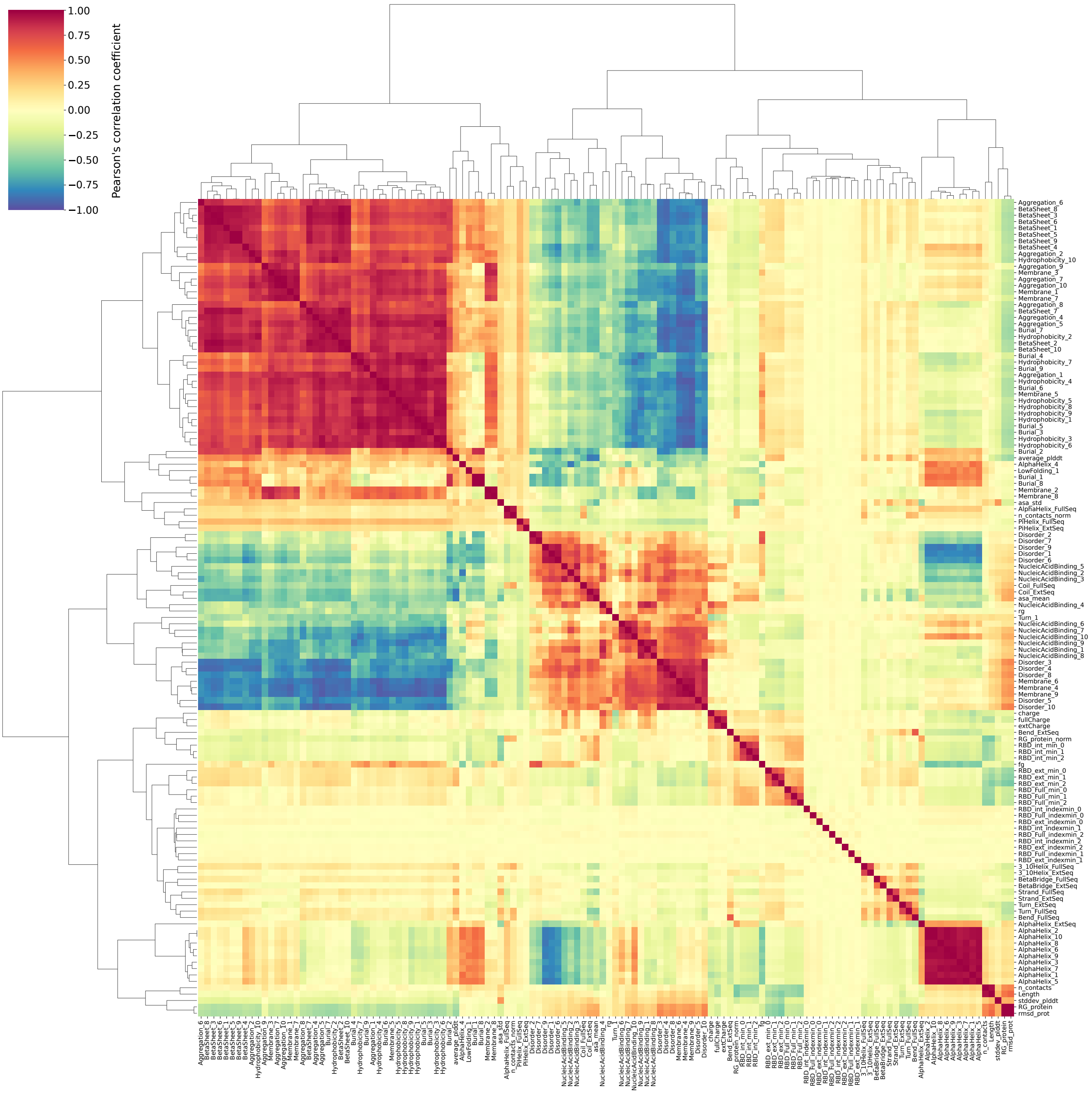


FIG. S3: Correlation cluster map of the 128 features used by catGRANULE 2.0 ROBOT to encode the proteins, computed from the training dataset. Refer to Supplementary Table S1 for the full feature names and their categorization into feature families.

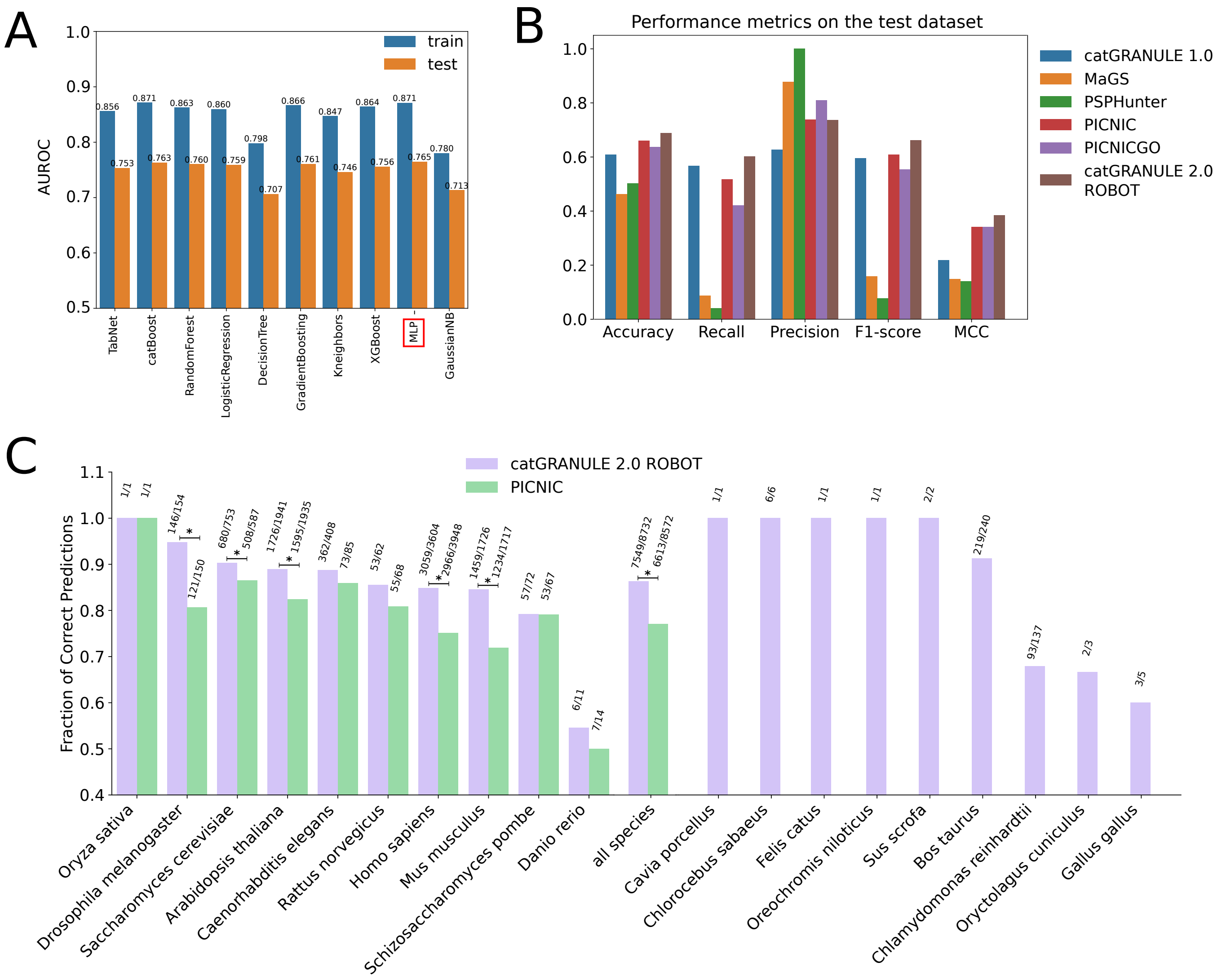


FIG. S4: A. AUROC score obtained by different classifiers on the training (blue) and test (orange) datasets. The selected model (MLP) is marked by a red square. B. Performance metrics of catGRANULE 2.0 ROBOT and other algorithms computed on the test dataset. MCC: Matthew's correlation coefficient. C. Bar plot of the fraction of correctly predicted LLPS proteins for different species. The annotation of LLPS proteins was obtained from the DrLLPS database [2]. A star above a bar indicates a p-value smaller than 0.05 from a Fisher's exact test between the fraction of correctly predicted LLPS proteins in catGRANULE 2.0 ROBOT and in PICNIC.

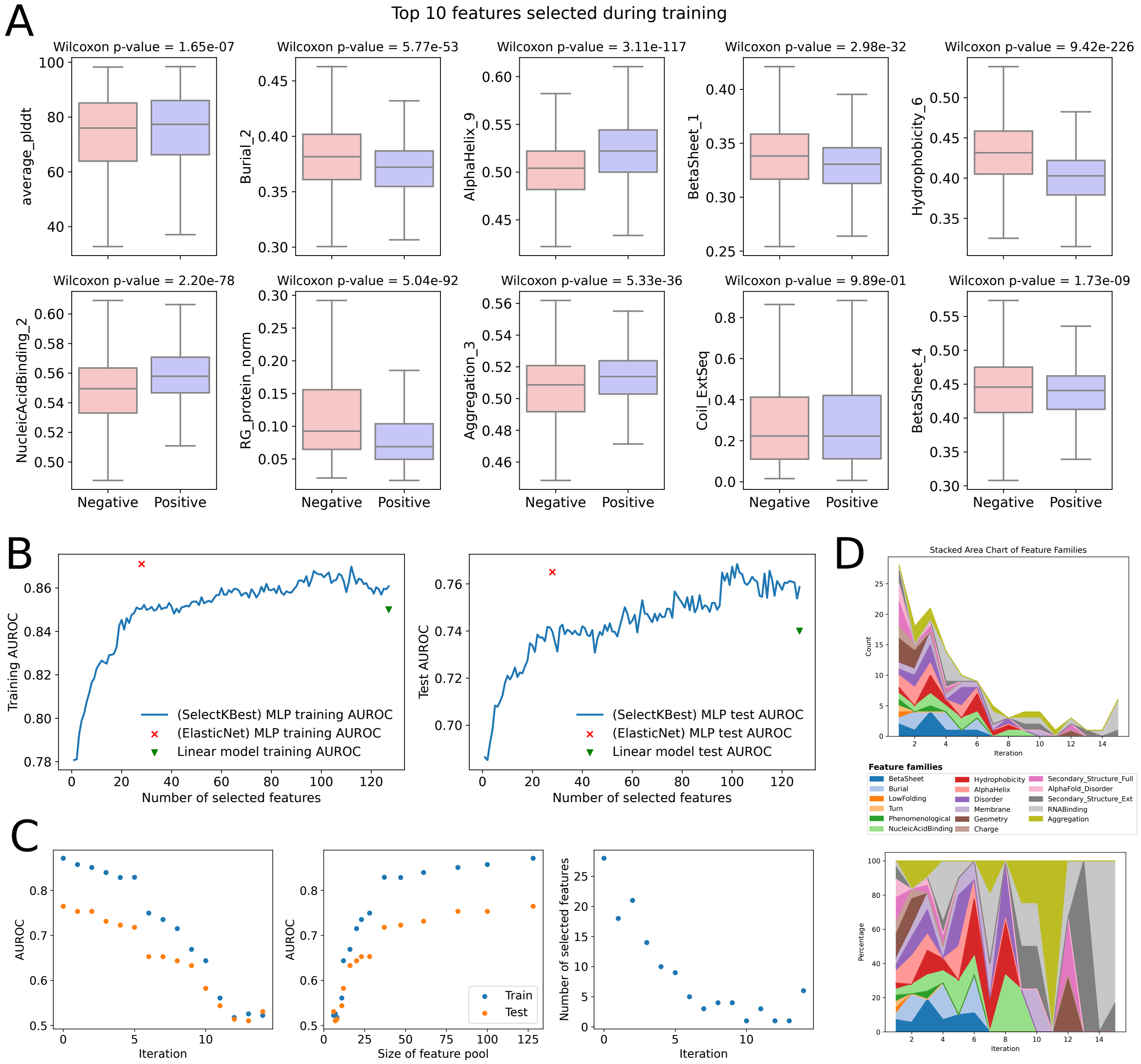


FIG. S5: A. Box plots of the values of the top 10 features, according to the permutation importance, selected during training, in the negative and positive training set. The p-value of the Wilcoxon rank-sum test is indicated in the title of each panel. B. AUROC score of a MLP classifier on the training (left) and test (right) datasets vs the number of features selected using SelectKBest, a univariate feature selection method. A green triangle indicates the AUROC of a linear model trained using the full set of features, the red x shows the AUROC achieved by selecting features with ElasticNet (the chosen model for catGRANULE 2.0 ROBOT). C. AUROC score of a MLP classifier on the training (blue) and test (orange) datasets obtained by removing the features selected by ElasticNet iteratively and repeating model training. The AUROC score is shown as a function of the number of iterations (left) and the size of the feature pool (middle), while the number of selected features versus the number of iterations is shown on the right. D. Stacked area chart of feature families vs iterations, shown by count (top) or by percentage (bottom) (refer to Supplementary Table S2 for the categorization of features into families).

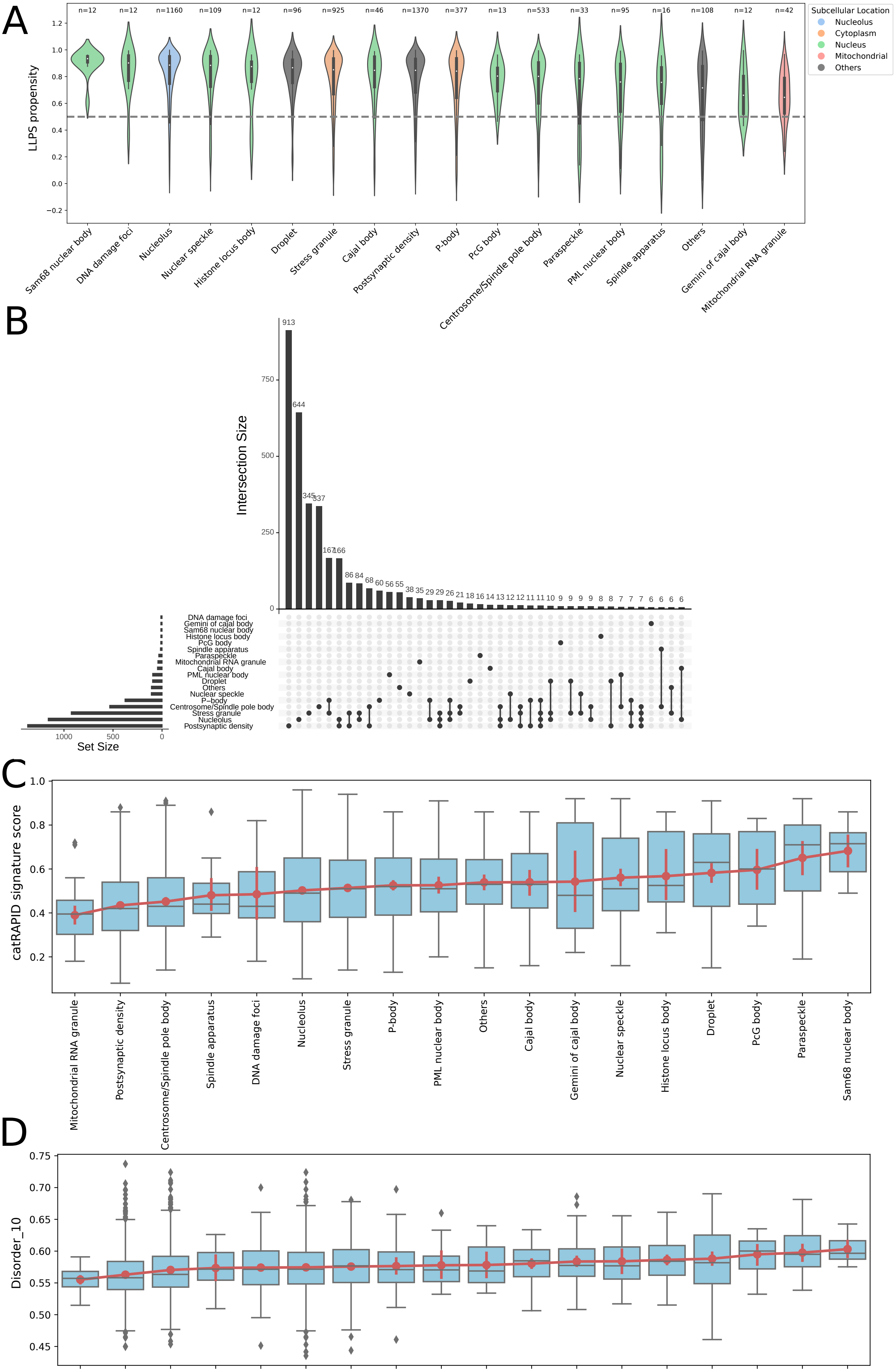
A
LLPS propensity
Subcellular Location
Nucleolus
Cytoplasm
Nucleus
Mitochondrial
Others
n=12
n=12
n=1160
n=109
n=12
n=96
n=925
n=46
n=1370
n=377
n=13
n=533
n=33
n=95
n=16
n=108
n=12
n=42
Sam68 nuclear body
DNA damage foci
Nucleolus
Nuclear speckle
Histone locus body
Droplet
Stress granule
Cajal body
Postsynaptic density
P-body
PcG body
Centrosome/Spindle pole body
Paraspeckle
PML nuclear body
Spindle apparatus
Others
Gemini of cajal body
Mitochondrial RNA granule
B
Intersection Size
Set Size
DNA damage foci
Gemini of cajal body
Sam68 nuclear body
Histone locus body
PcG body
Spindle apparatus
Paraspeckle
Mitochondrial RNA granule
Cajal body
PML nuclear body
Droplet
Others
Nuclear speckle
P−body
Centrosome/Spindle pole body
Stress granule
Nucleolus
Postsynaptic density
C
catRAPID signature score
Mitochondrial RNA granule
Postsynaptic density
Centrosome/Spindle pole body
Spindle apparatus
DNA damage foci
Nucleolus
Stress granule
P-body
PML nuclear body
Others
Cajal body
Gemini of cajal body
Nuclear speckle
Histone locus body
Droplet
PcG body
Paraspeckle
Sam68 nuclear body
D
Disorder_10

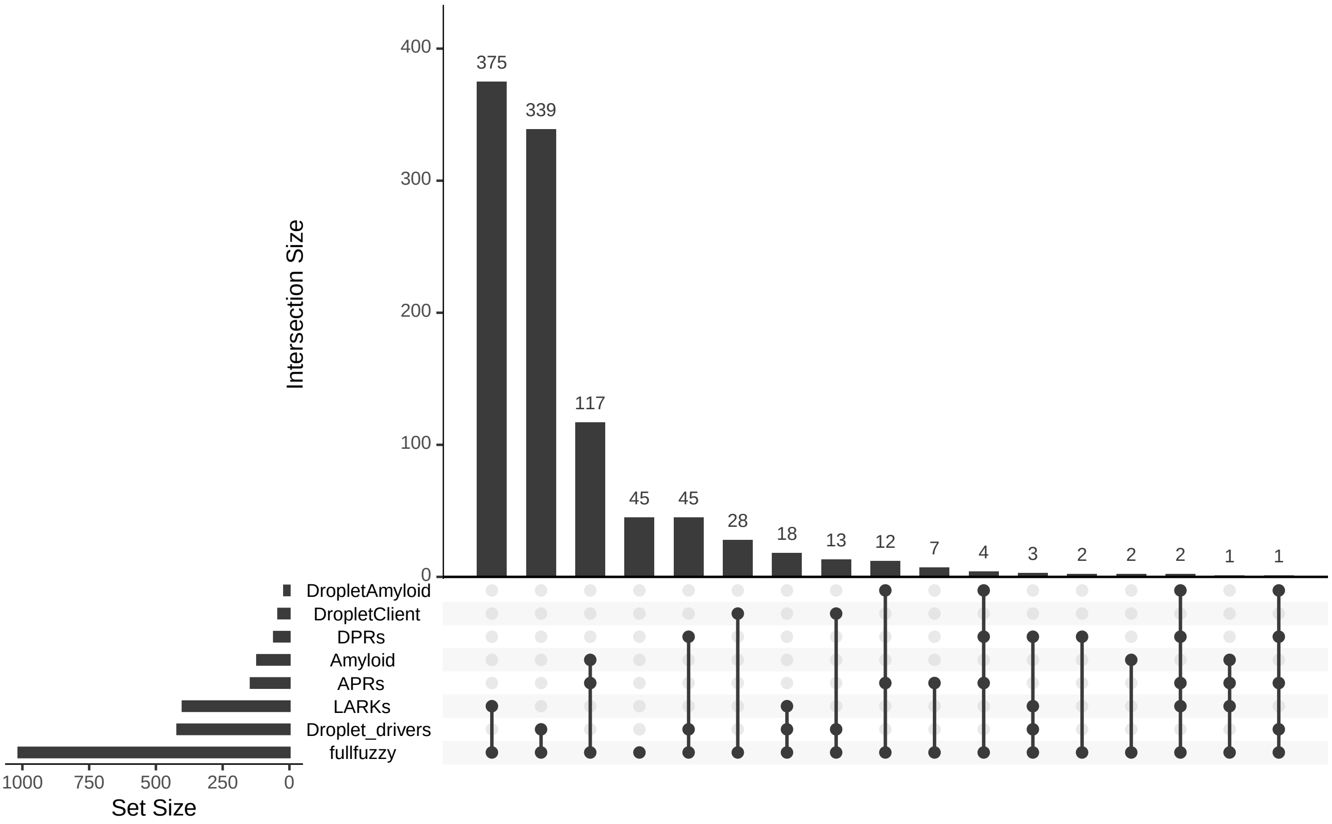


FIG. S7: Upset plot showing the number of proteins belonging to each category of condensate formation, defined in [? ], and the size of the intersections between different categories.

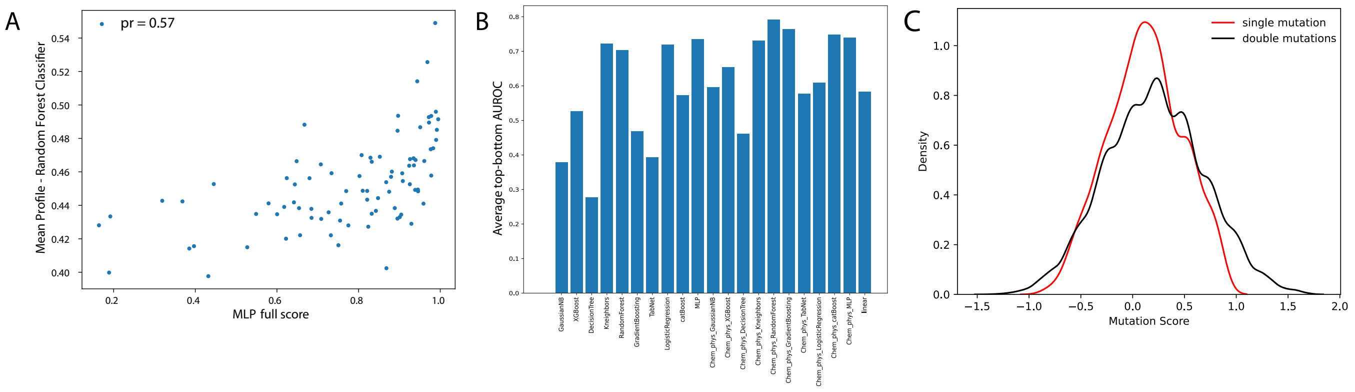


FIG. S8: A. Scatter plot of the LLPS propensity score of the MLP classifier trained using the full set of 128 features and the score obtained by averaging the LLPS propensity profile computed using a Random Forest classifier trained only on the set of physico-chemical features. The Pearson's correlation coefficient is reported in the figure legend. B. Average AUROC scores achieved by different classifiers, trained on the full set of features or only on the subset of physico-chemical features (indicated by the Chem_phys prefix in the x-axis tick labels), computed on the top and bottom scores based on experimentally annotated LLPS region of proteins collected from the PhaSepDB database (refer to Figure 5A and the Methods section). C. Distribution of the predicted catGRANULE 2.0 ROBOT mutation score for single and double amino acid mutations obtained from a mutational scanning of TDP-43. The distribution for the double mutations is wider than for the single, as expected, since cooperative mutations have a larger positive or negative effect on the LLPS propensity, compared to single amino acid mutations.
